# A gut-brain microbial sensing pathway mediates host defense and bacterial clearance

**DOI:** 10.64898/2026.10.07.757417

**Authors:** Alexander Runyon, Peizhao Zhang, Lihua Ye

## Abstract

The digestive tract is a major site of nutrient absorption. It also harbors the densest microbial communities in our body. It is well known that the brain perceives ingested nutrient information through the intestinal epithelium enteroendocrine cell (EEC)-vagal pathway. Little is known about how the brain senses gut bacteria that reside in the intestinal lumen. In this study, we combined novel in vivo EEC calcium imaging, EEC optochemical stimulation, and vagal sensory neuron calcium imaging in the zebrafish model, and revealed that segregated pathways mediated by different EEC and vagal sensory neuron subtypes differentially perceive beneficial nutrient information and harmful irritant pathogenic bacterial information. Our study revealed that a group of EECs that express the nociceptive receptor Transient receptor potential ankyrin A1 (Trpa1) can be activated by the pathogenic bacterium *Edwardsiella tarda* (*E. tarda*) or *E. tarda*-produced irritant metabolites. These Trpa1+EECs do not sense nutrients. Trpa1+EECs, but not other EECs, uniquely and highly express both Serotonin and the neuronal peptide Pituitary adenylate cyclase-activating polypeptide (Pacap). Using pharmaceutical and genetic approaches, our data showed that Pacap synergizes with Serotonin and is required for Trpa1+EEC-induced vagal activation. Stimulating the Trpa1+EEC-vagal sensory pathway potently inhibits a group of vagal motor neurons that innervate visceral organs in the caudal dorsal motor nucleus of the vagus (cDMV). However, the same group of cDMV neurons is activated by nutrients. Microbially induced cDMV inhibition enhances liver gluconeogenesis and promotes intestinal neutrophil recruitment. Inhibiting glucose metabolism, disrupting Trpa1+EEC-vagal sensory pathway, or optogenetically activating cDMV neurons in *E. tarda* colonized zebrafish induces intestinal *E. tarda* to switch from an aggregated phenotype to a diffused phenotype and hinders *E. tarda* clearance. Furthermore, our data show that disrupting the Trpa1+EEC-vagal sensory pathway impairs intestinal neutrophil recruitment and hinders survival, suggesting a protective role of the Trpa1+EEC-vagal microbial sensing pathway in pathogen defense. Together, our data revealed that a distinct non-nutrient-sensing Trpa1+EEC-vagal circuit senses gut microbial irritants, coordinating neuronal, metabolic, and immune responses to enhance host defense and pathogen clearance.

## Introduction

Trillions of diverse microbial species reside in the intestinal lumen. How the body senses and interacts with these gut microbial organisms is an exciting question that has received much attention in recent years. The intestinal mucosa forms a physical barrier that separates microbes from the body’s interior. Beyond the mucosal barrier, the body has evolved diverse systems that actively combat pathogen invasion while promoting tolerance to commensal microbiota^1,2^. To date, most research has focused on the protective role of the intestinal epithelial barrier and the immune system in defending against enteric pathogen invasion^3^. However, our understanding of other systems’ response to enteric microbial insults remains very limited.

The digestive tract is densely innervated by intrinsic and extrinsic neurons, which transmit diverse intestinal information to the central nervous system, enabling the brain to monitor the intestinal environment. Much is known about how the brain detects and responds to food and nutrient ingestion^4,5^. However, the means by which the brain detects and responds to commensal and pathogenic gut microbes remains largely unknown. As a result, our understanding of the central nervous system’s role in pathogen control and defense remains limited.

Recent studies have demonstrated that specific brain neurons can respond to systemic infection and inflammation^6–8^. It was shown that intraperitoneal injection of bacterial endotoxin Lipopolysaccharides (LPS) acutely activates a distinct group of neurons in the nucleus tractus solitarius (NTS) that express Adenylate Cyclase Activating Polypeptide 1 (Adcyap1), which drives sickness behavioral responses, including anorexia, decreased movement, and decreased core body temperature^7^. Bacterial infection-induced sickness responses, such as fever and anorexia, are believed to play an evolutionarily conserved protective role to enhance survival and promote immune defense and pathogen clearance^9–11^. The NTS receives sensory input from vagal sensory neurons and is the brain’s portal for integrating visceral information. Recent research demonstrated that distinct vagal sensory neurons encode systemic LPS induced inflammation signals, which may drive neuronal activation in the NTS^8^. These recent studies provide valuable insights into the critical role of neuronal responses to systemic infection, a process that occurs once pathogenic bacteria pass the intestinal mucosal barrier. In addition to systemic infection, earlier studies have shown that the central nervous system also responds to local enteric bacterial infection^12,13^. For example, it was shown that gavaging the enteric pathogen Campylobacter jejuni to the mouse intestine acutely activates NTS neurons in the absence of elevated levels of circulating pro-inflammatory cytokines^12^, suggesting that the brain may also be able to detect and respond to pathogens that are confined to the intestinal lumen and have not yet passed the mucosal barrier. However, how the brain detects pathogenic bacteria in the intestinal lumen remains largely unknown. Whether and how the brain coordinates the physiological and immune response to fight enteric pathogen invasion is also unclear.

The intestinal epithelium is the frontline of the mucosa that directly interacts with the intestinal lumen microbial components. Within the intestinal epithelium reside a group of specialized sensory cells called enteroendocrine cells (EECs). EECs exhibit huge heterogeneity and sense diverse information in the intestinal lumen^14,15^. Much is known about EECs’ ability to sense macro- and micro-nutrients related to food ingestion^14,15^. Recent studies have demonstrated that certain EECs may also sense microbially derived signals^16–19^. For example, it has been shown that EECs that secrete serotonin (enterochromaffin cells or ECs) can sense microbially derived isovalerate through olfactory receptor 558^17,18^. Isovalerate-induced EC activation induces visceral hypersensitivity through the spinal sensory nerve^17,18^. Using the zebrafish model, our previous studies also demonstrated that a subset of EECs that express Transient Receptor Potential Ankyrin 1 (Trpa1) can be activated by fish pathogenic bacteria *Edwardsiella tarda* (*E. tarda*)^19^. However, these Trpa1-expressing EECs are not activated by commensal fish bacterial species^19^. We further demonstrated that *E. tarda* secreted tryptophan metabolites such as indole or indole-3-aldehyde (I3A) are novel Trpa1 agonists that potently activate Trpa1-expressing EECs^19^. However, it remains unknown why Trpa1+EECs sense *E. tarda* or *E. tarda* derived microbial irritant signals, and how Trpa1+EECs transmit the microbial signals.

In this study, using in vivo calcium imaging together with genetic manipulation, we demonstrated a comprehensive pathway by which Trpa1+EECs communicate with a subset of vagal sensory neurons to transmit intestinal microbial irritant signals to the brain and coordinate systemic defense responses to eliminate invading enteric pathogens and protect the host.

## Results

### Microbial irritants, but not nutrients, activate zebrafish Trpa1+EECs

Trpa1 is expressed in a subset of EECs, and these Trpa1-expressing EECs are distributed along the intestine (**Fig. 1A**). To further understand the genetic profile of these Trpa1-expressing EECs (hereafter Trpa1+EECs), we utilized a previously published single-cell RNA seq dataset^20^. Our analysis demonstrated that Trpa1+EECs are segregated from the other EEC subtypes (**Fig. 1B-D, Fig. S1A-E**). Trpa1+EECs do not express colonic nutrient sensory EEC markers such as Cholecystokinin (CCK), Peptide YY (PYY), or Glucagon-like peptide 1 (GLP-1) (**Fig. 1B-D, Fig. S1A-E**), indicating that Trpa1+EECs have a unique genetic profile compared with other EECs.

**Figure 1.**
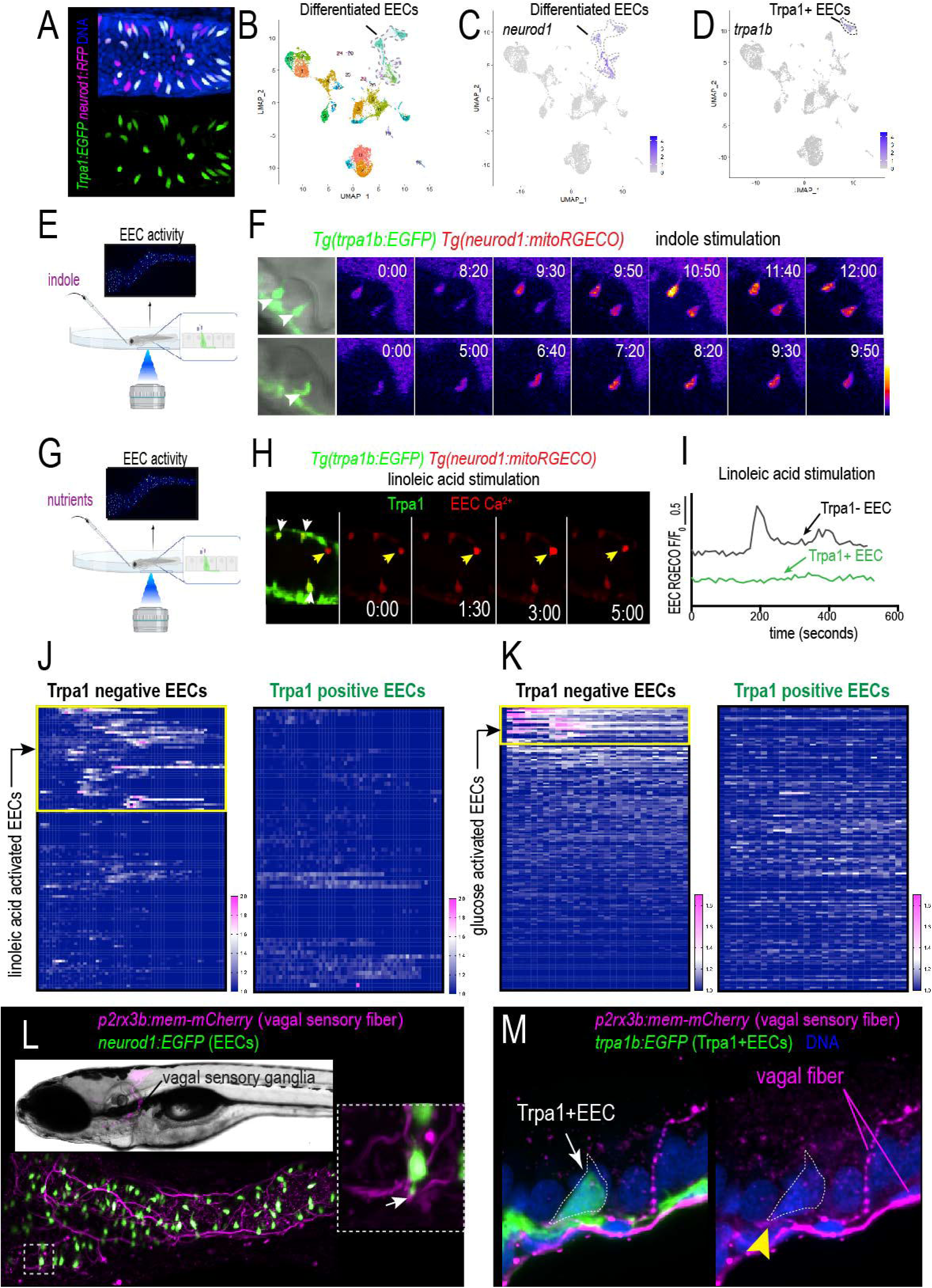
Trpa1+EECs are activated by indole but not nutrients. **(A)** Representative confocal image shows Trpa1-expressing EECs in the intestinal epithelium. EECs are labeled by *Tg(neurod1:RFP)*, and Trpa1+EECs are labeled by *Tg(trpa1b:EGFP)*. Trpa1+EECs are indicated by white arrows. **(B-D)** Zebrafish intestinal epithelium single-cell RNA sequencing dataset from Wen J. et al., 2021, showing that Trpa1+EECs form a distinct cell cluster separate from other EECs. **(E-F)** In vivo EEC Ca^2+^ imaging showing indole activates Trpa1+EECs. Trpa1+EECs are labeled by *Tg(trpa1b:EGFP)* (white arrows in F), and EEC Ca^2+^ activity is labeled by *Tg(neurod1:mitoRGECO)*. **(F)** Time-lapse image frames showing increased mitoRGECO fluorescence in three representative Trpa1+EECs post indole stimulation. **(G-K)** In vivo EEC Ca^2+^ imaging showing fatty acid and glucose do not activate Trpa1-expressing EECs. **(H)** Time-lapse image frames showing mitoRGECO fluorescence in four EECs post linoleic acid stimulation. The three Trpa1+EECs are labeled by *Tg(trpa1b:EGFP)* (white arrows), and one EECs that does not express Trpa1 are labeled by a yellow arrow. Note that the EECs that do not express Trpa1 are activated by linoleic acid, and the three nearby Trpa1+EECs are not activated. **(I)** Normalized mitoRGECO fluorescence post linoleic acid stimulation in representative Trpa1- and Trpa1+ EECs. **(J-K)** Clustering analysis of Trpa1- and Trpa1+ EECs Ca^2+^ activity following linoleic acid or glucose stimulation. 71 Trpa1+EECs and 114 Trpa1-EECs from 5 fish were analyzed for the linoleic acid response. 152 Trpa1+EECs and 167 Trpa1-EECs from 4 fish were analyzed for the glucose response. EECs activated by linoleic acid or glucose are boxed in yellow. Note that only Trpa1-EECs are activated by nutrients. **(L)** Top: Representative *Tg(p2rx3b:mem-mCherry)* transgenic zebrafish, in which the vagal sensory neurons are labeled with magenta. Bottom: Live imaging of a zebrafish intestine showing EECs and vagal sensory fibers. Dashed box shows an EEC connecting with the vagal sensory fiber. **(M)** Confocal image showing a Trpa1+EEC directly contacting a vagal sensory fiber (yellow arrowhead).

To further analyze the sensory capacity of the Trpa1+EECs, we used an EEC in vivo calcium imaging approach^21^ that we previously developed (**Fig. S2**). Similar to mammals, nutrients like glucose and fatty acids activate a subset of EECs and increase cytoplasmic Ca^2+^ (**Fig. S2**). The elevated cytoplasmic Ca^2+^ flux into mitochondria triggers mitochondrial Ca^2+^ increase^21^. This can be measured using the *Tg(neurod1:mitoRGECO)* genetic model, which expresses a red-fluorescent calcium indicator in EEC mitochondria^21^. We then crossed *Tg(trpa1b:EGFP)* with *Tg(neurod1:mitoRGECO)*, in which Trpa1+EECs can be identified by green fluorescence EGFP and EEC Ca2+ can be measured via red fluorescence RGECO. First, we stimulated zebrafish with indole, a metabolite secreted by *E. tarda* that activates Trpa1+EECs. We found that indole activates Trpa1+EECs and increases mitoRGECO fluorescence signal (**Fig. 1E-F**). Our previous and current results showed that nutrients such as linoleic acid (a long-chain fatty acid) or glucose stimulation activate a subset of EECs (**Fig. S2**)^21,22^. Next, we stimulated zebrafish with linoleic acid and glucose. Interestingly, our results showed that linoleic acid or glucose activate a subset of EECs that do not express Trpa1, and Trpa1+EECs are not activated by these nutrients (**Fig. 1G-I**). Consistent with this finding, single-cell RNA sequencing analysis also showed that Trpa1-expressing EECs do not express glucose or fatty acid sensors (**Fig. S1F-G**). Together, these results demonstrated that Trpa1+EECs are unique sensory EECs in the intestinal epithelium. These cells exhibit a distinct genetic profile that is separate from other nutrient-sensing EECs. Importantly, our in vivo EEC activity recording demonstrated that these Trpa1+EECs exhibit unique sensory capacity, in that they are activated by microbial irritants such as indole but not nutrients.

### Stimulating Trpa1+EECs activates a subtype of vagal sensory neurons

Previous studies have shown that nutrient-sensing EECs (such as CCK+EECs) communicate with vagal sensory neurons, and a subset of EECs form synaptic-like connections with vagal sensory nerve fibers in the intestine^23^. We recently developed a *Tg(p2rx3b:mem-mCherry)* transgenic zebrafish line that labels vagal sensory neurons^24^. Like mammals, some zebrafish EECs directly connect with vagal sensory fibers (**Fig. 1L**). Using the *Tg(p2rx3b:mem-mCherry)* model, our data showed that a subset of Trpa1+EECs also form direct connections with vagal sensory nerve fibers (**Fig. 1M**), suggesting that EEC-vagal connection and communication may not be restricted to nutrient sensing EECs.

To further investigate whether and how Trpa1+EECs communicate with vagal sensory neurons, we used our previously established optochemical approach to stimulate Trpa1+EECs via light^19^ and imaged the vagal sensory neurons’ calcium activity in live zebrafish. In this approach, zebrafish were treated with optovin, which binds the Trpa1 channel and activates the Trpa1 channel in a UV light-dependent manner (**video1**). Using a customized confocal setup (see Materials and Methods), we can deliver 405nm light specifically to the zebrafish intestine region and image the vagal sensory neurons’ calcium activity simultaneously (**Fig. 2A-C**). First, we tested whether we can precisely deliver light in a region-specific manner. To do this, we used a *Tg(neurod1:mitoEOS)* zebrafish model, in which the photoconvertible EOS protein is expressed in EECs, pancreatic islets, and neurons. We defined the region of interest (ROI) in the zebrafish midgut and delivered 405nm light to the defined ROI (**Fig. S3**). Our results showed that 405nm light only converts the EOS protein from green to red in the defined ROI, and other regions remain unconverted (**Fig. S3**). Next, we performed the experiment in optovin-treated zebrafish. 405nm light is delivered to the middle intestine region to activate Trpa1+EECs, and vagal sensory neuron calcium activity is measured via *Tg(elavl3:H2BGcamp6s)* (**Fig. 2B**). *Tg(elavl3:H2BGcamp6s)* expresses a genetically encoded calcium indicator in neuronal nuclei and is widely used for zebrafish neuronal calcium imaging. Our data revealed that stimulating Trpa1+EECs in the intestine activates a subset of vagal sensory neurons (**Fig. 2D-K**) (**Video 2**). The vagal sensory neuron calcium activity returns to the basal level when 405nm light is off (**Fig. 2D-K**) (**Video 2**), indicating that the observed vagal sensory neuron activation is dependent on Trpa1+EEC stimulation. Repeated Trpa1+EEC stimulation via 405nm light activates the same subset of vagal sensory neurons, although the calcium amplitude is lower compared to the initial activation (**Fig. 2J**). Consistently, in a *Tg(neurod1:Gcamp6f)* model that measures vagal sensory neuron cytoplasmic calcium activity, we also observed that stimulating Trpa1+EECs activates a subset of vagal sensory neurons (**Fig. S4A-G**). Vagal sensory neurons are not activated when 405nm light is delivered to zebrafish intestine that is not treated with Optovin (**Fig. S4H-J**), suggesting that the observed vagal sensory neuron activation is triggered by Trpa1+EEC stimulation but not 405nm light. Together, our data suggest that zebrafish Trpa1+EECs communicate with vagal sensory neurons, and stimulating Trpa1+EECs activates a subset of these neurons.

**Figure 2.**
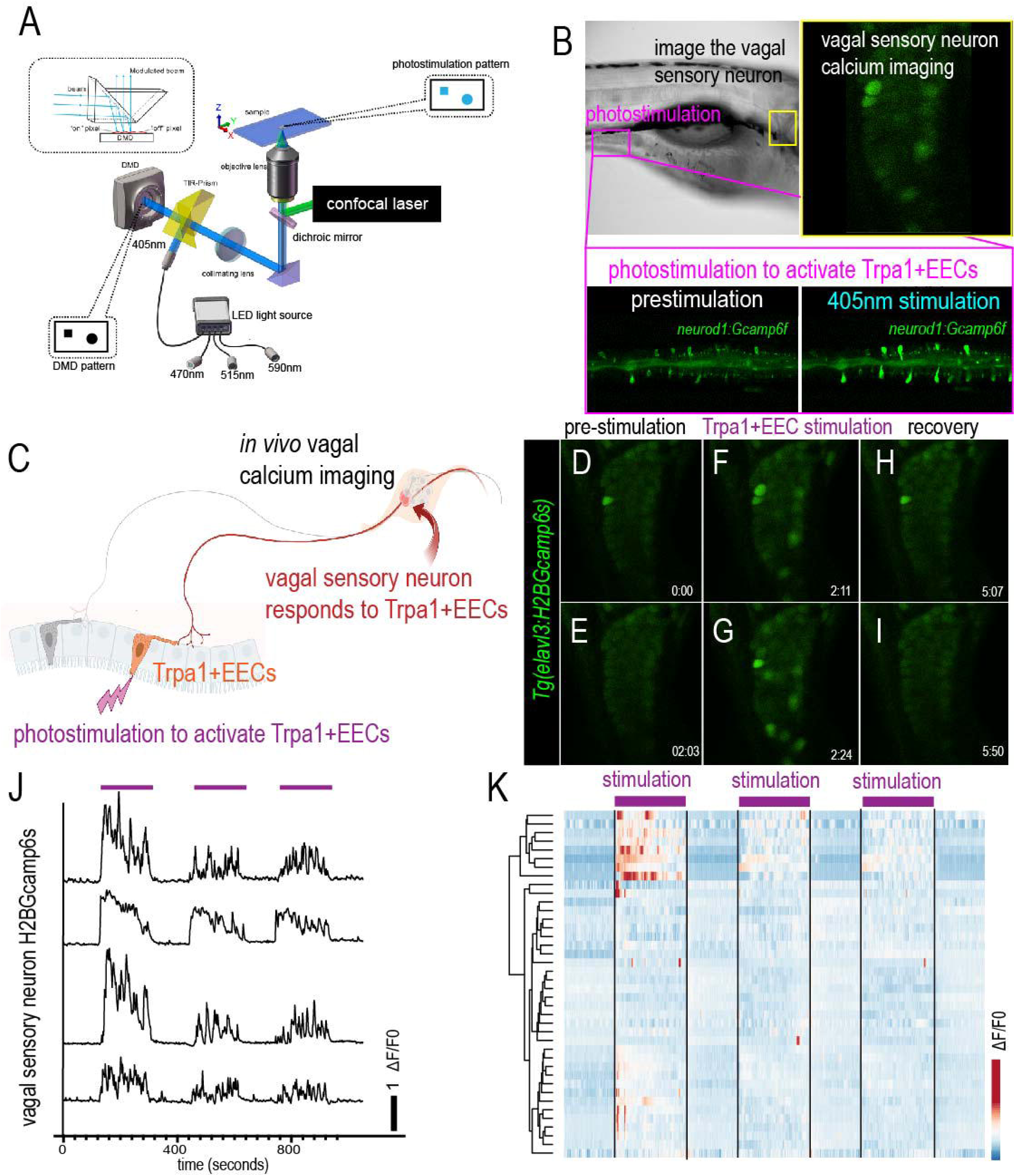
Stimulating Trpa1+EECs activates a subset of vagal sensory neurons. **(A-C)** Equipment setup and schematic model showing simultaneous light stimulation to activate Trpa1+EECs and image vagal sensory neuron calcium activity. The stimulating light at a specific wavelength was delivered via an LED light source separate from the confocal imaging laser. A digital micromirror device (DMD) was used to control the specificity of the light stimulation region. Zebrafish were treated with Optovin, and 405nm light was delivered to the intestinal region to activate Trpa1+EECs. EEC activity was labeled by *Tg(neurod1:Gcamp6f)*, and vagal activity was labeled by *Tg(elavl3:H2BGcamp6s)*. The light-stimulating region and the vagal imaging region are boxed in B. **(D-I)** Representative vagal sensory neuron calcium activity before, during, and after Trpa1+EEC light stimulation. Stimulating Trpa1+EECs activates a subset of vagal sensory neurons. **(J)** Plot of four representative vagal sensory neuron calcium profiles in response to Trpa1+EEC stimulation. Purple lines indicate the time when 405nm light was on to stimulate Trpa1+EECs. Vagal sensory neuron H2BGcamp6s was normalized to t_0_, and ΔF/F0 was plotted. **(K)** Clustering analysis and heatmap of vagal sensory neurons’ calcium activity. 40 vagal sensory neurons from 3 fish were analyzed.

Previous mammalian studies showed that nutrients such as fatty acids or glucose activate vagal sensory neurons^25,26^. We then investigated how zebrafish vagal sensory neurons respond to nutrient stimulation. We stimulated zebrafish with linoleic acid and glucose and monitored the vagal sensory neuron calcium activity via *Tg(elavl3:H2B-Gcamp6s)*. Our data revealed that in response to linoleic acid stimulation, a subset of vagal sensory neurons is activated (**Fig. 3A-D**). Similarly, when stimulating with glucose, we observed that glucose also activates a subset of vagal sensory neurons (**Fig. S5A, B, E**) (**Video 3**). We did not detect vagal sensory neuron activation in response to water stimulation (**Fig. S5C-D**). Together, these data suggest that, like mammals, zebrafish vagal sensory neurons also respond to nutrient stimuli.

**Figure 3.**
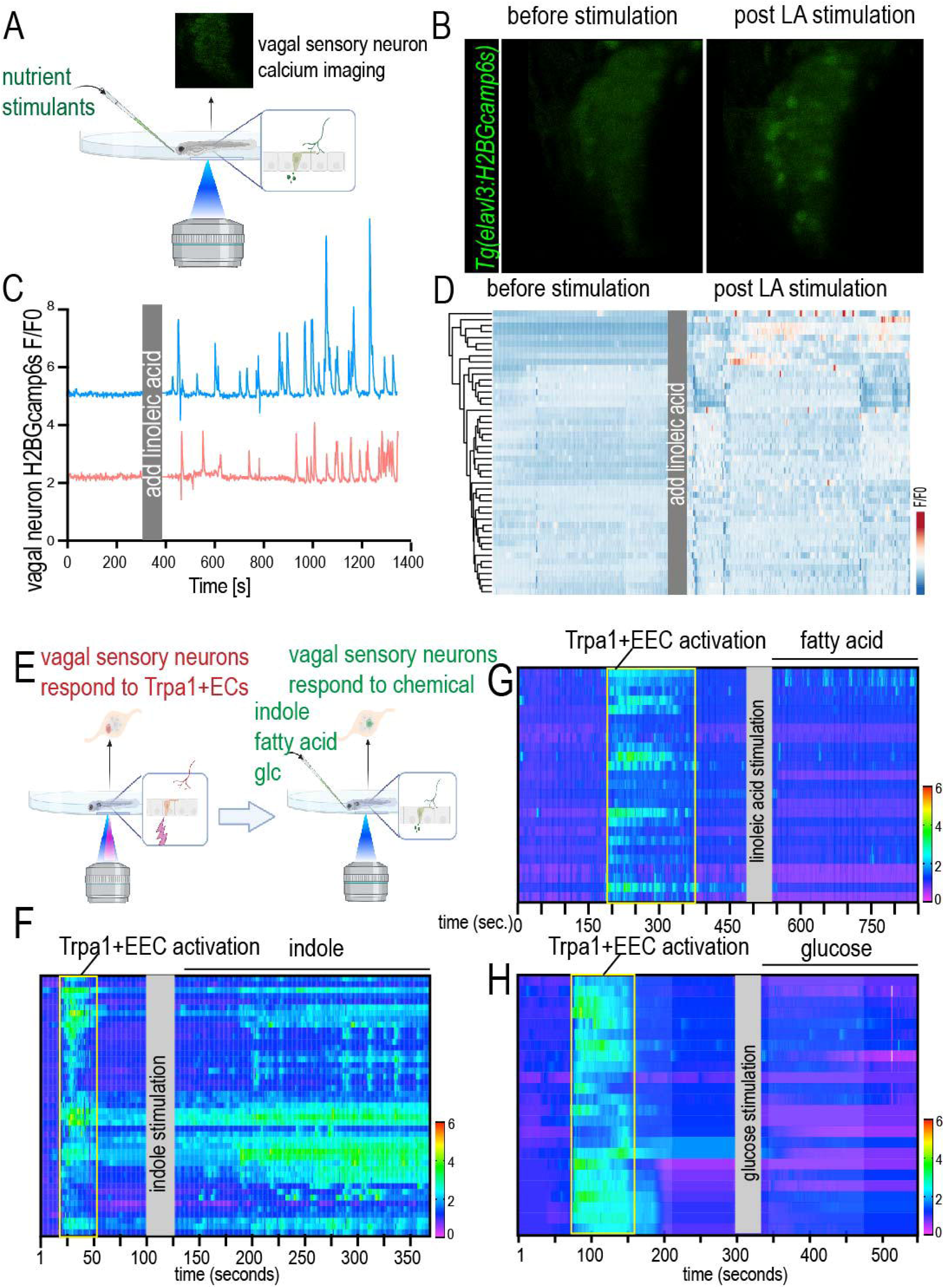
Trpa1+EEC-activated vagal sensory neurons do not sense nutrients. **(A-D)** Imaging vagal sensory neuron calcium activity in response to nutrient stimulation. **(A)** Experimental setup. **(B)** Representative vagal sensory ganglion calcium activity before and 5 minutes post linoleic acid stimulation. Vagal sensory neuron calcium activity was measured via *Tg(elavl3:H2BGcamp6s)*. **(C)** Two representative linoleic acid-activated vagal sensory neuron calcium activity plots before and after linoleic acid stimulation. **(D)** Clustering analysis and heatmap of vagal sensory calcium activity before and after linoleic acid stimulation in one representative zebrafish. Note that a subset of vagal sensory neurons was activated in response to linoleic acid stimulation. **(E)** Experimental setup showing vagal sensory neuron calcium imaging in response to sequential optic stimulation to activate Trpa1+EECs and chemical stimulants. 405nm light was delivered to the intestine in Optivin-treated zebrafish, and vagal sensory neuron activity was recorded. Different chemical stimulants (indole, linoleic acid, and glucose) were then delivered to the same zebrafish, and calcium activity in the same group of vagal sensory neurons was analyzed. **(F)** Clustering and heatmap plot showing that vagal sensory neurons activated by Trpa1+EEC stimulation are activated by indole. 45 Trpa1+EEC activated vagal sensory neurons were analyzed. **(G-H)** Clustering and heatmap plot showing that most of the vagal sensory neurons activated by Trpa1+EEC stimulation are not activated by linoleic acid or glucose. 25 Trpa1+EEC-activated vagal sensory neurons were analyzed in the linoleic acid-stimulating group. 24 Trpa1+EEC-activated vagal sensory neurons were analyzed in the glucose-stimulating group.

In light of this, we investigated whether Trpa1+EECs and nutrients activate the same group of vagal sensory neurons. We stimulated Trpa1+EECs via the optochemical approach as described above and imaged the vagal sensory neuron calcium via *Tg(elavl3:H2BGcamp6s)*. The vagal sensory neuron calcium activity was recorded before, during, and after Trpa1+EEC stimulation. After vagal sensory neuron calcium activity returned to baseline, we stimulated the same zebrafish with different chemical stimulants, including indole, linoleic acid, and glucose. Our previous study and the data above demonstrated that indole stimulates Trpa1+EECs (**Fig. 1E-F**)^19^. Our results showed that the vagal sensory neurons that are activated by Trpa1+EECs also respond to indole stimulation as expected (**Fig. 3F**). However, the vagal sensory neurons that are activated by Trpa1+EECs do not respond to fatty acid or glucose stimulation (**Fig. 3G, H**). Together, our data suggest that nutrients and bacterial irritants such as *E. tarda* or *E. tarda* secreted metabolites are perceived by different EECs. *E. tarda* or *E. tarda*-secreted indole activates Trpa1+EECs. Nutrients such as fatty acids or glucose activate another group of EECs that do not express Trpa1. These different EEC subgroups communicate to different vagal sensory neurons, and the irritant bacterial information transmitted by Trpa1+EECs is sensed by a distinct group of vagal sensory neurons that do not sense nutrients.

### Serotonin and Pacap are involved in Trpa1+EEC-vagal communication

After demonstrating that stimulating Trpa1+EECs activates a subgroup of vagal sensory neurons, we then investigated how Trpa1+EECs communicate with vagal sensory neurons. Our previous study^19^ and new analysis of the recent scRNA-sequencing dataset revealed that Trpa1+EECs but not other EECs or IECs express *tph1b*, a key enzyme for serotonin synthesis (**Fig. 4A-B**). In addition to Serotonin, our data analysis also showed that Trpa1+EECs highly express *adenylate cyclase activating polypeptide 1a* (*adcyap1a*), which encodes the neuronal peptide Pituitary adenylate cyclase-activating polypeptide (Pacap) (**Fig. 4C**). This expression is unique to Trpa1+EECs as other EECs or IECs do not express *adcyap1a* or Pacap (**Fig. 4C**). The expression of Serotonin and Pacap in Trpa1+EECs were further confirmed via immunofluorescence staining (**Fig. 4D-E**). We then questioned whether Trpa1+EECs communicate with vagal sensory neurons via Serotonin or Pacap. To answer this question, we first searched a published zebrafish larvae scRNA-seq dataset^27^ to examine whether serotonin or Pacap receptors are expressed in the zebrafish vagal sensory neurons. Our analysis revealed that htr3a, but not other serotonin receptors, is highly expressed in vagal sensory neurons (**Fig. S6**). Htr3a is not expressed in enteric neurons, and its expression in other CNS neurons is very low (**Fig. S6A**). The expression of htr3a in vagal sensory neurons is further confirmed using Hybridization Chain Reaction In Situ Hybridization (HCR-ISH) (**Fig. 4F**). Similarly, our results showed that zebrafish vagal sensory neurons express the Pacap receptor adcyap1r1a (**Fig. 4G**, **Fig. S6B**); however, unlike htr3a, *adcyap1r1a* is broadly expressed in peripheral and central neurons (**Fig. S6B**).

**Figure 4.**
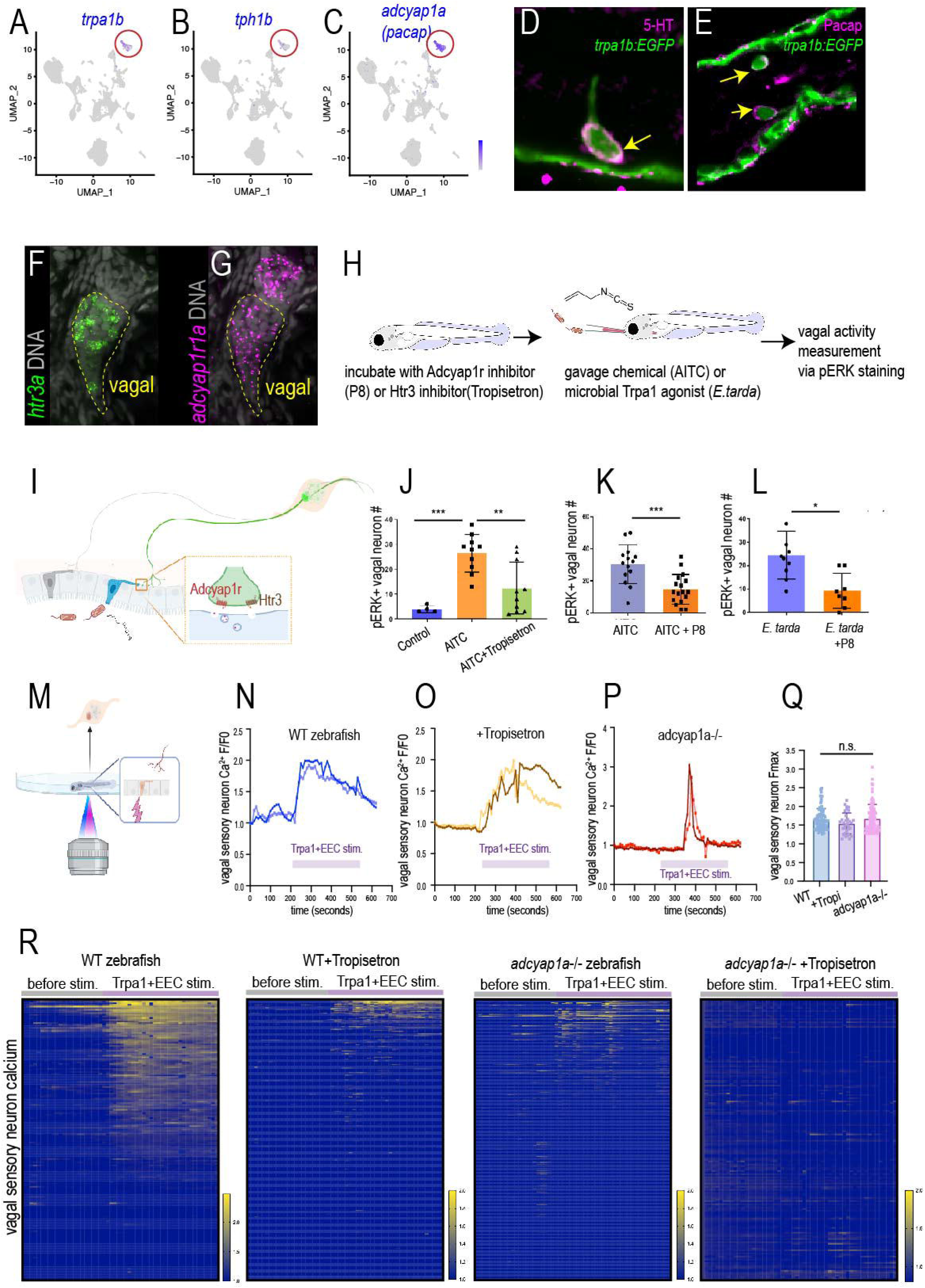
Trpa1+EECs communicate with vagal sensory neurons via serotonin and PACAP. **(A-C)** UMAP plot of *trpa1b*, *tph1b*, and *adcyap1a* genes from the zebrafish intestinal epithelium single-cell RNA sequencing dataset by Wen J. et al., 2021. Note that only Trpa1+EECs, but not other EECs or IECs, express *tph1b* or *adcyap1a* (*pacap*). **(D-E)** Confocal images of immunofluorescence staining showing that Trpa1+EECs express serotonin and Pacap. Trpa1+EECs were labeled by *Tg(trpa1b:EGFP)*. **(F-G)** HCR-ISH images showing that zebrafish vagal sensory neurons express htr3a and adcyap1r1a. **(H-I)** Experimental setup and schematic model to investigate the role of serotonin and PACAP signaling in Trpa1+EEC-vagal communication. Zebrafish were pretreated with HTR3 inhibitor Tropisetron or Pacap receptor inhibitor P8. Following HTR3 or Pacap receptor inhibition, zebrafish were gavaged with TrpA1 agonist AITC or *E.tarda*. Vagal sensory neuron calcium activity was analyzed via pERK staining. **(J-L)** Quantification of pERK+ vagal sensory neuron number in different experimental groups. **(J)** Tropisetron inhibits AITC induced vagal sensory neuron activation. **(K-L)** P8 inhibits AITC or *E.tarda* induced vagal sensory neuron activation. **(M-R)** Image vagal sensory neuron calcium activity in response to Trpa1+EEC optic stimulation. Vagal sensory neuron calcium activity was measured by *Tg(elavl3:H2BGcamp6s)*. **(N-P)** Representative activated vagal sensory neuron calcium plot in response to Trpa1+EEC stimulation in WT, tropisetron-treated, and *adcyap1a*-/-zebrafish. Note that vagal sensory neurons exhibit different activation profiles in response to Trpa1+EEC stimulation. **(Q)** Activated vagal sensory neuron maximum H2BGcamp6s fluorescence in response to Trpa1+EEC stimulation. **(R)** Heatmap plots of vagal sensory neuron calcium profile before and post Trpa1+EEC stimulation in WT, Tropisetron-treated, *adcyap1a*-/-, and Tropisetron-treated *adcyap1a*-/- zebrafish. Note that both Tropisetron treatment and *adcyap1a*-/- reduce the number of vagal sensory neurons that are activated in response to Trpa1+EEC stimulation. WT: 171 vagal sensory neurons from 5 zebrafish. Tropisetron: 311 vagal sensory neurons from 9 zebrafish. *adcyap1a*-/-: 435 vagal sensory neurons from 9 zebrafish. *adcyap1a*-/- +Tropisetron: 177 vagal sensory neurons from 5 zebrafish. J, K: One-way ANOVA with Tukey posttest. K, L: Student’s t-test. * P<0.05, ** P<0.01, *** P<0.001.

Next, we used the pharmacological approach to investigate whether serotonin or Pacap signaling is involved in Trpa1+EEC-vagal communication. To this end, we pretreated zebrafish with the highly selective HTR3a inhibitor Tropisetron or the Pacap receptor inhibitor P8^28–30^. We then gavaged Trpa1 agonist Allyl isothiocyanate (AITC) or *E. tarda* into the zebrafish larvae’s intestine and examined vagal sensory neuron activity via pERK immunofluorescence staining^19,31^. Our results demonstrated that inhibition of HTR3a or Pacap receptor reduced AITC- or *E. tarda*-induced vagal activation (**Fig. 4H-L**), suggesting that serotonin and Pacap signaling may both be involved in Trpa1+EEC-vagal communication.

To further investigate the role of serotonin signaling in mediating Trpa1+EEC-vagal signaling transmission, we pretreated zebrafish with Tropisetron and performed in vivo vagal sensory calcium imaging using *Tg(elavl3:H2BGcamp6s)*. Our data revealed that inhibiting HTR3a reduced the number but not the excitation amplitude of the vagal sensory neurons in response to Trpa1+EEC stimulation (**Fig. 4M-O, Q, R**), suggesting that serotonin signaling is required for Trpa1+EEC-vagal communication.

Pacap receptor is broadly expressed in the central and peripheral nervous system. To further investigate the role of Pacap signaling in mediating Trpa1+EECs-induced vagal activation and downstream physiology, we used the CRISPR/cas9 approach to knock out the *adcyap1a* gene and generated *adcyap1a*-/- zebrafish (**Fig. S7**). Zebrafish have two adcyap1 orthologs: *adcyap1a* and *adcyap1b*. Data from whole zebrafish scRNA sequencing demonstrated that *adcyap1a* and *adcyap1b* exhibit different expression patterns. Adcyap1a is highly and specifically expressed in EECs, while *adcyap1b* is expressed in the nervous system but not EECs (**Fig. S7**). Therefore, mutating *adcyap1a* will affect Pacap signaling specifically in Trpa1+EECs but not in the nervous system. Consistent with this, immunofluorescence staining also revealed that in WT zebrafish, Pacap is expressed in both EECs and enteric neurons (**Fig. S7B**). In *adcyap1a*-/- zebrafish, the expression of Pacap in EECs is eliminated, while Pacap expression in enteric neurons remains intact (**Fig. S7C**). Using this model, we then tested the role of Trpa1+EEC Pacap signaling in mediating Trpa1+EEC-vagal communication. Our data showed that in WT zebrafish, vagal sensory neurons exhibit sustained activation during the Trpa1+EEC stimulation phase (**Fig. 2**, **Fig. 4N**). Loss of Pacap in Trpa1+EECs significantly reduced the number of vagal sensory neurons that were activated by Trpa1+EECs (**Fig. 4R**). Moreover, these activated vagal sensory neurons exhibit a transient activation peak rather than sustained activation during the Trpa1+EEC stimulation phase (**Fig. 4N, P**). The amplitude of vagal sensory neuron activation is not reduced in *adcyap1a*-/- zebrafish (**Fig. 4Q**). When we inhibit HTR3a via Tropisetron in *adcyap1a*-/- zebrafish, Trpa1+EEC induced vagal sensory neuron activation is nearly abolished (**Fig. 4R**). Together, these data suggest that both Serotonin and Pacap signaling are required to mediate Trpa1+EEC-vagal communication. Serotonin and Pacap signaling exhibit synergistic effects in sustaining vagal sensory neuron activation in response to Trpa1+EEC stimulation.

### Stimulating Trpa1+EECs inhibits caudal DMV neurons

Our data demonstrated that stimulating Trpa1+EECs activates a subgroup of vagal sensory neurons, and that Trpa1+EEC-vagal communication is mediated by serotonin and Pacap signaling. Next, we questioned how Trpa1+EECs affect brain activity. Like mammals, zebrafish vagal sensory neurons send signals to the nucleus tractus solitarius (NTS) in the hindbrain^24^. NTS neurons process vagal sensory information and relay the signal to vagal motor neurons in the dorsal motor nucleus of the vagus (DMV) (**Fig. 5A**) (**Video 5**). Zebrafish vagal motor neurons innervate visceral organs and are critical in regulating physiological homeostasis^32,33^. We then investigated how different stimuli regulate zebrafish DMV neuron activity. Vagal motor neurons’ calcium activity was measured via *Tg(chata:Gal4); Tg(UAS:Gcamp6s)*. Our data revealed in unstimulated zebrafish, most DMV neurons exhibit low GCaMP activity, while a subgroup of vagal motor neurons in the caudal DMV exhibits high basal GCaMP activity (**Fig. S8**). When zebrafish are stimulated with nutrients such as glucose or linoleic acid, this group of caudal DMV neurons increases activity (**Fig. 5B-D**) (**Video 6**). Interestingly, when zebrafish were stimulated with indole, we observed strong inhibition of caudal DMV neurons, while some neurons in the rostral and middle DMV regions were activated (**Fig. 5M-S**) (**Video 7**). To further investigate how Trpa1+EECs modulate DMV neurons, we used the optochemical approach above to stimulate Trpa1+EECs via light and imaged DMV neuron calcium activity. Consistently, our data revealed that stimulating Trpa1+EECs acutely activates rostral and middle DMV neurons but inhibits a subset of caudal DMV neurons (**Fig. 5T-V**). We then treated zebrafish with indole for 12 hours and examined DMV neuron Gcamp activity. We observed that 12h-indole treatment reduces caudal but not rostral DMV neuronal activity (**Fig. S9A-D**). Using a gnotobiotic approach, we colonized germ-free (GF) zebrafish with *E.tarda* bacteria or commensal microbiota and analyzed DMV neuron activity via pERK staining. Our data showed that *E. tarda,* but not commensal microbiota colonization, inhibits caudal DMV neuron activity (**Fig. S9E-H**).

**Figure 5.**
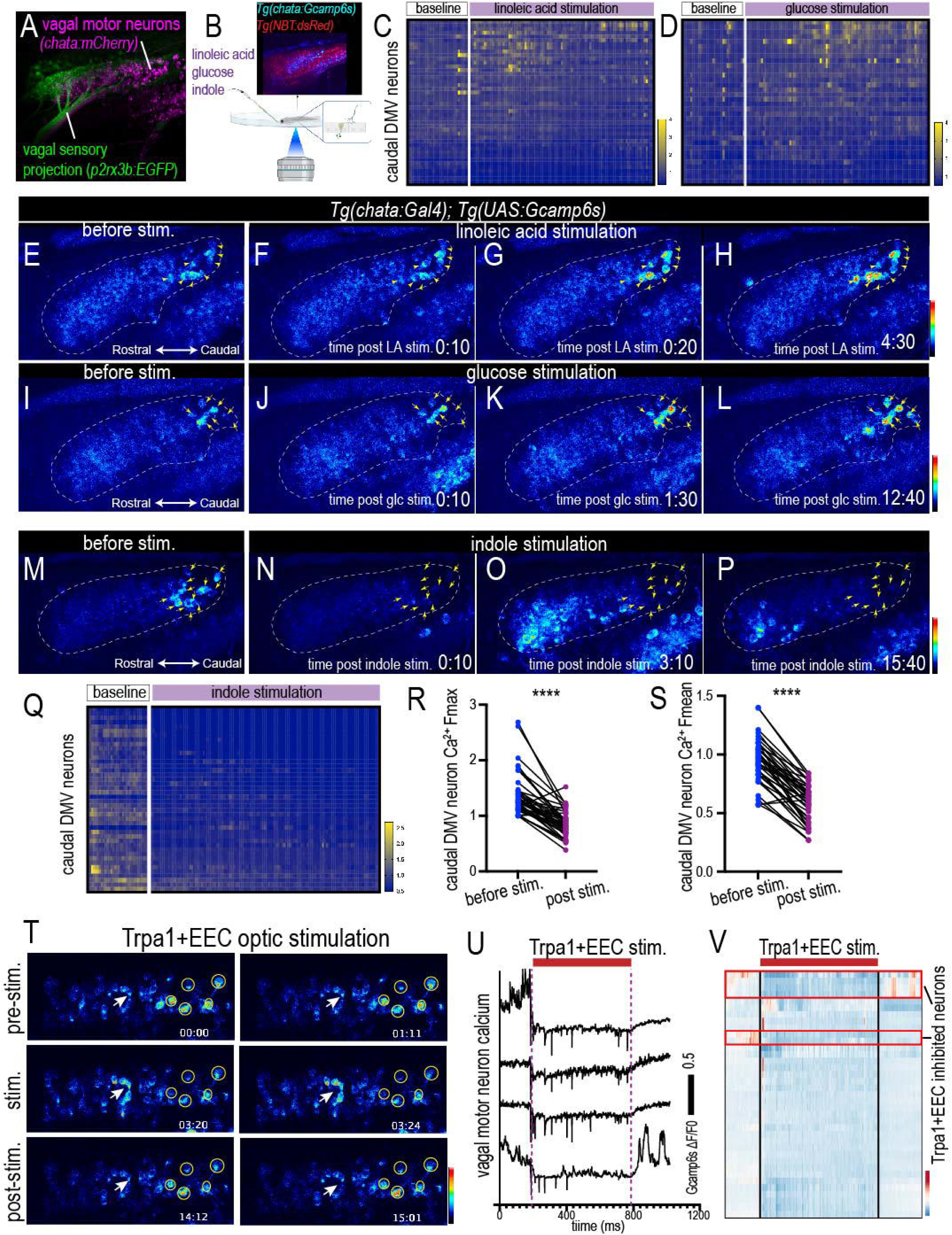
Stimulating Trpa1+EECs inhibits caudal DMV neuron activity. **(A)** Confocal imaging showing vagal sensory neuron central projection and DMV vagal motor neurons in 7 dpf zebrafish hindbrain. The vagal sensory neuron central projection was labeled by *Tg(p2rx3b:EGFP)*, and vagal motor neurons were labeled by *Tg(chata:Gal4); Tg(UAS:NTR-mCherry)*. **(B-S)** Imaging vagal motor neuron calcium activity in response to chemical stimulants. The vagal motor neuron calcium activity was measured via *Tg(chata:Gal4); Tg(UAS:Gcamp6s)*. 300 seconds were imaged before stimulation, and 900 seconds were imaged post-stimulation in the glucose and linoleic acid stimulation groups. For indole stimulation group, 1200 seconds were imaged post stimulation. **(B)** Experimental setup. **(C-D)** Heatmap plots of caudal DMV neuron calcium profile in response to linoleic acid or glucose stimulation. Caudal DMV neuron Gcamp fluorescence was normalized to T_0_. **(E-P)** Representative DMV neuron calcium activity before and after linoleic acid, glucose, or indole stimulation. Note that in basal conditions, a subset of vagal motor neurons exhibits higher calcium activity compared to the remaining DMV. Linoleic acid or glucose enhances activity in a subset of high-activity caudal DMV neurons (yellow arrowheads in E-L), while indole potently inhibits these caudal DMV neurons (yellow arrowheads in M-P). **(Q)** Heatmap plot of caudal DMV neurons before and after indole stimulation. Caudal DMV neuron Gcamp fluorescence was normalized to T_0_. **(R-S)** Maximum and mean caudal DMV neuron Gcamp fluorescence before and after indole stimulation. Each dot in R and S represents an individual vagal motor neuron. 44 caudal DMV neurons from 4 zebrafish were analyzed in C. 34 caudal DMV neurons from 4 zebrafish were analyzed in D. 42 caudal DMV neurons from 6 zebrafish were analyzed in Q-S. **(T-V)** Imaging vagal motor neuron calcium activity in response to Trpa1+EEC optic stimulation. Vagal motor neuron calcium was measured by *Tg(chata:Gal4); Tg(UAS:Gcamp6s)*. Trpa1+EECs were stimulated by 405nm light in Optovin-treated zebrafish. **(T)** Representative DMV vagal motor neuron calcium before, during, and after Trpa1+EEC stimulation. White arrows: DMV neurons that are activated by Trpa1+EEC stimulation. Yellow arrows: caudal DMV neurons that are inhibited by Trpa1+EEC stimulation. **(U)** Four representative caudal DMV neurons that are inhibited by Trpa1+EEC stimulation. **(V)** Clustering and heatmap showing DMV neuron calcium activity before, during, and after Trpa1+EEC stimulation. Red box: Trpa1+EEC inhibited vagal motor neuron. 37 DMV neurons were analyzed. R, S: Wilcoxon paired tests. **** P<0.0001.

Together, our data show that DMV neurons are differentially modulated by nutrients and microbial irritants. Specifically, our data showed that nutrients such as glucose or fatty acids excite a subset of cDMV neurons that exhibit high basal activity. However, these same groups of cDMV neurons are potently inhibited by microbial irritants that activated Trpa1+EECs.

### Stimulating Trpa1+EECs with bacterial irritants increases liver gluconeogenesis

Previous studies in zebrafish demonstrated that DMV neurons are anatomically organized along the rostral-caudal axis^32,34^, with rostral and middle DMV neurons innervating the pharyngeal arch while caudal DMV neurons innervate visceral organs such as the intestine and liver^32,34^. Stimulating cDMV neurons modulates visceral organ calcium activity, suggesting a conserved feedback vagal motor neuron in zebrafish larvae^33^. Vagal motor neurons are critical for regulating liver physiology, and inhibiting vagal motor neurons enhances liver gluconeogenesis^35–38^. Liver gluconeogenesis is controlled by the rate-limiting enzyme phosphoenolpyruvate carboxykinase 1 (Pck1)^39^. Pck1 transcription is tightly controlled by hormonal and neuronal signaling, including cholinergic signaling from vagal motor neurons^36,39–41^. To investigate how *E. tarda* bacteria or indole affects liver gluconeogenesis, we used a *Tg(pck1:venus, cryaa:RFP)* zebrafish reporter line^42^. Our data show that treating zebrafish with indole for 12 hours significantly elevated liver *pck1:venus* fluorescence (**Fig. 6A-E**). Consistently, indole treatment increased zebrafish systemic glucose levels (**Fig. 6G**). In addition to indole, I3A, the other *E. tarda* tryptophan metabolite that activates Trpa1+EECs, also increased liver gluconeogenesis (**Fig. 6F**). However, tryptophan metabolites that do not activate Trpa1+EECs do not alter liver gluconeogenesis (**Fig. 6H**), suggesting that indole and I3A induced liver gluconeogenesis may be associated with their ability to activate Trpa1+EECs. As a control experiment, we also found that feeding reduces liver gluconeogenesis as expected (**Fig. 6H**). To further examine how *E. tarda* bacteria affect liver gluconeogenesis, we colonized GF zebrafish with *E. tarda* bacteria or commensal microbiota. Our data showed that *E. tarda* colonization promotes gluconeogenesis, while commensal bacteria colonization did not alter liver gluconeogenesis (**Fig. 6I**).

**Figure 6.**
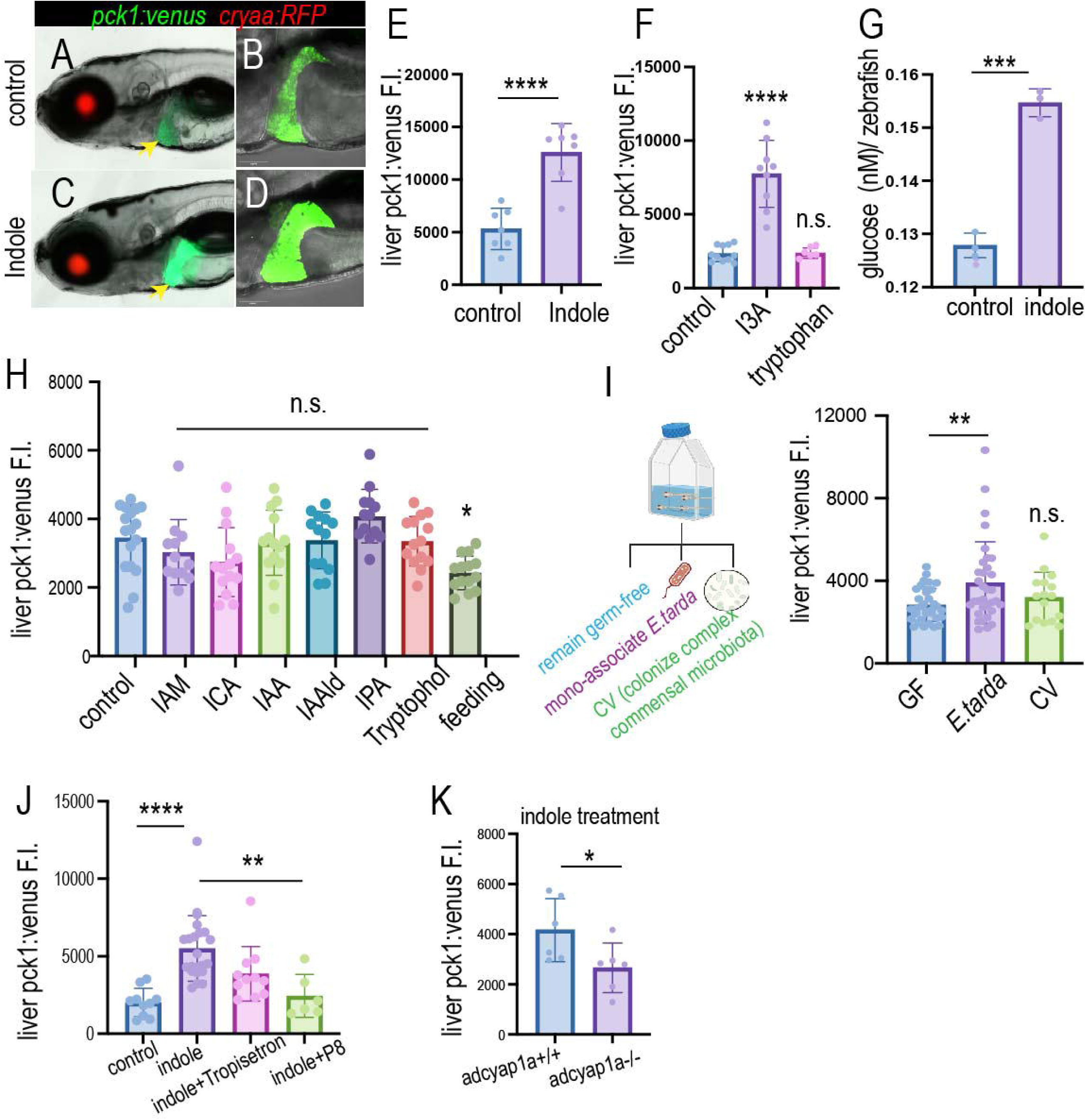
Stimulating Trpa1+EECs via microbial irritants increases liver gluconeogenesis. **(A-D)** Epifluorescence (A, C) and confocal (B, D) images of control and 12-hour indole-treated *Tg(pck1:venus, cryaa:RFP)* zebrafish. **(E-F)** Quantification of liver pck1:venus fluorescence intensity in control, indole, I3A, or Tryptophan treated zebrafish. **(G)** Quantification of free glucose level in control and indole-treated zebrafish. **(H)** Quantification of liver *pck1:venus* fluorescence intensity in zebrafish that were treated with different tryptophan metabolites for 12 hours or in zebrafish fed with larvae food for 12 hours. I3A: indole-3-aldehyde; IAM: Indole-3-acetamide; ICA: Indole-3-carboxylic acid; IAA: Indole-3-acetic acid; IAAld: Indole-3-acetaldehyde; IPA: Indole-3-pyruvic acid. **(I)** Gnotobiotic approach to Monoassociate germ-free zebrafish with *E. tarda* or Commensal microbiota from 3 dpf to 7 dpf. Liver pck1:venus fluorescence was analyzed in 7 dpf zebrafish. GF: Germ-free. CV: Commensal microbiota colonized conventionalized. **(J)** Quantification of liver *pck1:venus* fluorescence in control or indole-treated zebrafish without or with Tropisetron or P8. (K) Quantification of liver pck1:venus fluorescence in WT or *adcyap1a*-/- zebrafish after 12 hours of indole treatment.

We then investigated the role of Trpa1+EEC Serotonin and Pacap signaling in indole-induced liver gluconeogenesis increase. We treated zebrafish with HTR3a inhibitor Tropisetron or Pacap receptor inhibitor P8. Our data showed that P8, but not Tropisetron, significantly reduced the indole-induced increase in liver gluconeogenesis (**Fig. 6J**). Consistently, our data also showed that loss of Trpa1+EEC Pacap signaling in *adcyap1a*-/- zebrafish decreased liver gluconeogenesis in response to indole treatment (**Fig. 6K**). Together, these data suggest that indole-induced liver gluconeogenesis requires Trpa1+EEC Pacap signaling.

Data from zebrafish larvae single-cell RNA sequencing and our HCR-ISH showed that zebrafish larvae liver does not express the Pacap receptor (**Fig. S10**), suggesting that Pacap released from Trpa1+EECs may indirectly regulate liver gluconeogenesis rather than act directly on hepatocytes. Because Trpa1+EEC Pacap signaling plays an important role in sustaining vagal sensory neuron activation, as shown above, we questioned whether indole induces liver gluconeogenesis through the Trpa1+EEC-vagal axis. To answer this question, we developed a zebrafish model to ablate vagal sensory neurons. In this model, a genetically modified bacterial Nitroreductase (NTR_2.0_) gene is expressed in the vagal sensory neurons via the Gal4/UAS system^43^. Expression of the NTR_2.0_ gene in vagal sensory neurons converts the prodrug Metronidazole (MTZ) into a toxic form, thereby ablating these neurons (**Fig. 7A-D**). We used this model to ablate vagal sensory neurons. The transgenic negative zebrafish that received the same MTZ treatment were used as a control. Following vagal ablation, we treated zebrafish without or with indole. Our results showed that in control zebrafish, indole significantly increases liver gluconeogenesis (**Fig. 7E-F**). However, indole induced liver gluconeogenesis increase is significantly impaired in vagal ablated zebrafish (**Fig. 7E-F**), suggesting that vagal sensory neurons are required for indole induced liver gluconeogenesis increase.

**Figure 7.**
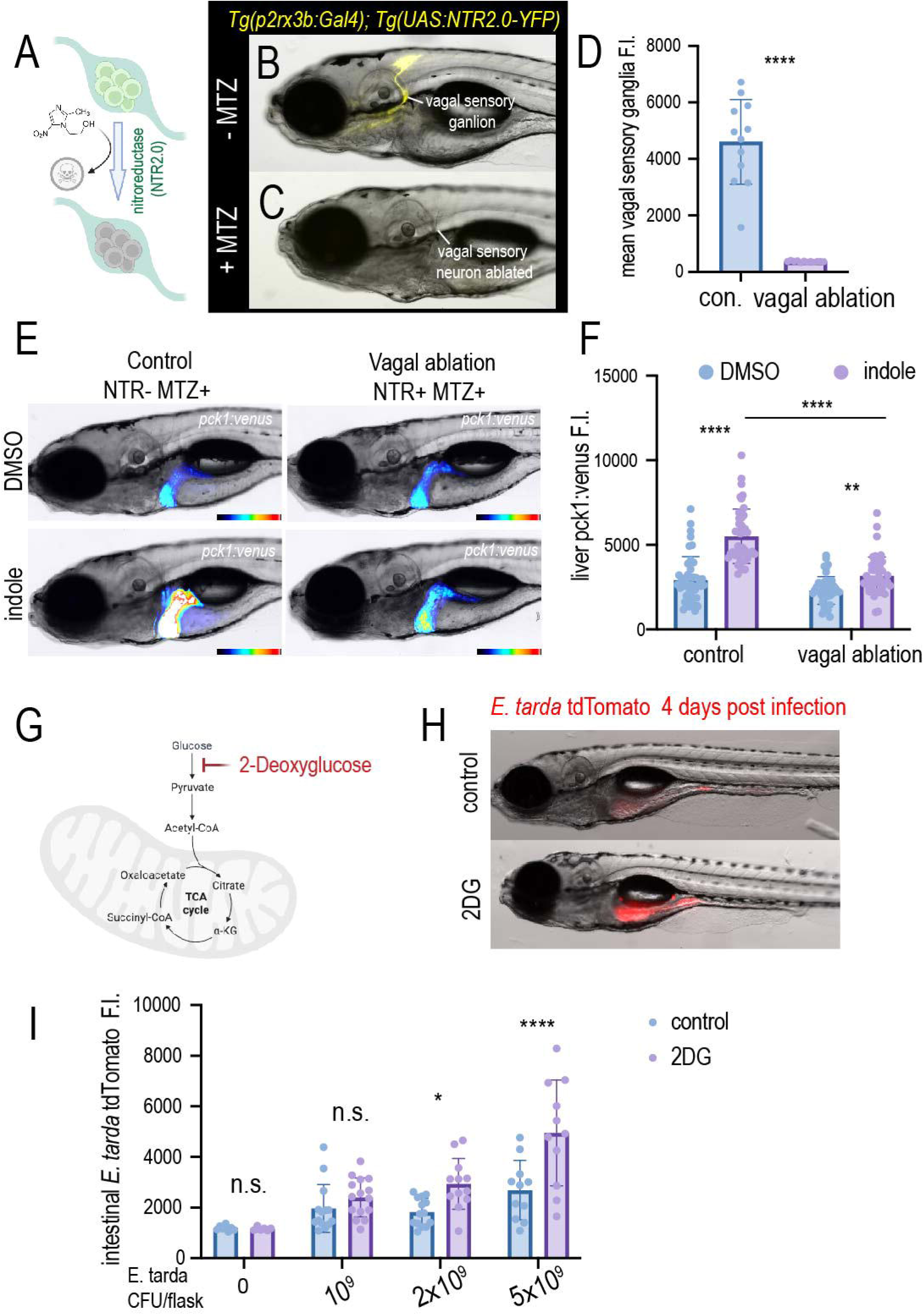
Indole-induced liver gluconeogenesis requires vagal sensory neurons. **(A-D)** A new genetic model to ablate vagal sensory neurons in zebrafish. (A) Schematic of vagal sensory neuron ablation via the Nitroreductase (NTR) system. NTR_2.0_-YFP transgene was specifically expressed in vagal sensory neurons via the *Tg(p2rx3b:Gal4); Tg(UAS:NTR2.0-YFP)* transgenic model. NTR_2.0_ converts the prodrug Metronidazole (MTZ), which is not harmful to zebrafish cells, into a toxic form and kills the cells. (B-C) Representative *Tg(p2rx3b:Gal4); Tg(UAS:NTR2.0-YFP)* zebrafish without or with MTZ treatment. Note that the YFP+ vagal sensory neurons were completely ablated by MTZ. (D) Quantification of NTR_2.0_-YFP fluorescence in the vagal sensory ganglion in zebrafish without or with MTZ treatment. **(E)** Epifluorescence images of *Tg(pck1:venus)* control and vagal ablated zebrafish without or with indole treatment. **(F)** Quantification of liver pck1:venus fluorescence intensity in control and vagal ablated zebrafish without or with indole treatment. **(G)** Schematic model of how 2-deoxyglucose (2DG) inhibits glucose metabolism. **(H)** Representative image of zebrafish that were treated without or with 2DG and infected with *E. tarda*. Zebrafish were infected with different doses of *E. tarda* tdTomato from 3 dpf to 7 dpf and were imaged at 7 dpf. **(I)** Quantification of intestinal *E. tarda* fluorescence intensity in control and 2DG-treated zebrafish. D: Student’s t-test. F, I: Two-way ANOVA with Tukey posttest. ** P<0.01, **** P<0.0001.

### Dysfunction of the Trpa1+EEC-vagal pathway induces *E. tarda* diffusion and impairs *E. tarda* clearance

Our data showed that stimulating Trpa1+EECs with microbial irritants increases liver gluconeogenesis and systemic glucose via Trpa1+EEC PACAP signaling and the vagal sensory neuron pathway. We then examined how systemic glucose increases and glucose metabolism affect host defense against irritant bacteria such as *E. tarda*. To answer this question, we developed an *E. tarda* bacterial strain that stably expresses tdTomato fluorescence using the mini-Tn7 transposon approach^44^. We then treated zebrafish with 2-Deoxy-D-glucose (2-DG), a non-metabolizable glucose analog that inhibits glycolysis and glucose metabolism^45^. We infected control or 2-DG treated zebrafish with different doses of *E. tarda*. Our data showed that inhibiting glucose metabolism with 2-DG impairs *E. tarda* clearance from the intestine and increases intestinal lumen *E. tarda* abundance (**Fig. 7G-I**), suggesting that systemic glucose increase may play a beneficial role in response to acute enteric bacterial insults.

Because Trpa1+EEC PACAP signaling and vagal sensory neurons are involved in microbial irritant-induced liver gluconeogenesis and systemic glucose increase, we then investigated the role of Trpa1+EEC PACAP signaling and vagal sensory neurons in *E. tarda* clearance. To this end, we infected WT and *adcyap1a*-/- zebrafish with different doses of *E.tarda* tdTomato. Our results showed that in WT zebrafish, *E. tarda* typically clustered in large aggregates (**Fig. 8A**, **Fig. S11A**). This is true in both high- and low-dose *E. tarda* colonization conditions (**Fig. 8A**, **Fig. S11A**). Interestingly, in *adcyap1a*-/- zebrafish, *E. tarda* was dispersed in the intestinal lumen, and *E. tarda* tdTomato fluorescence in the intestine increased significantly (**Fig. 8A-B**, **Fig. S11A**). We then performed confocal imaging of WT and *adcyap1a*-/- zebrafish that were infected with *E. tarda* tdTomato. We observed that the large *E. tarda* aggregates in WT zebrafish do not touch the intestinal epithelium and are not motile (**Fig. 8C-D**) (**Video 8**). In *adcyap1a*-/- zebrafish, *E. tarda* are dispersed as individual *E. tarda* cells or very small clusters (**Fig. 8E-H**). Some of the small *E.tarda* clusters or *E. tarda* cells directly touch the intestinal epithelium (**Fig. 8E-H**). However, we did not detect *E. tarda* inside intestinal epithelial cells in WT or *adcyap1a*-/- zebrafish. The dispersed *E. tarda* cells or *E. tarda* small clusters in *adcyap1a*-/- zebrafish intestine are motile (**Video 9**). These data suggest that loss of Pacap in Trpa1+EECs induces *E. tarda* to switch from an aggregated, non-motile form to a dispersed, motile form.

**Figure 8.**
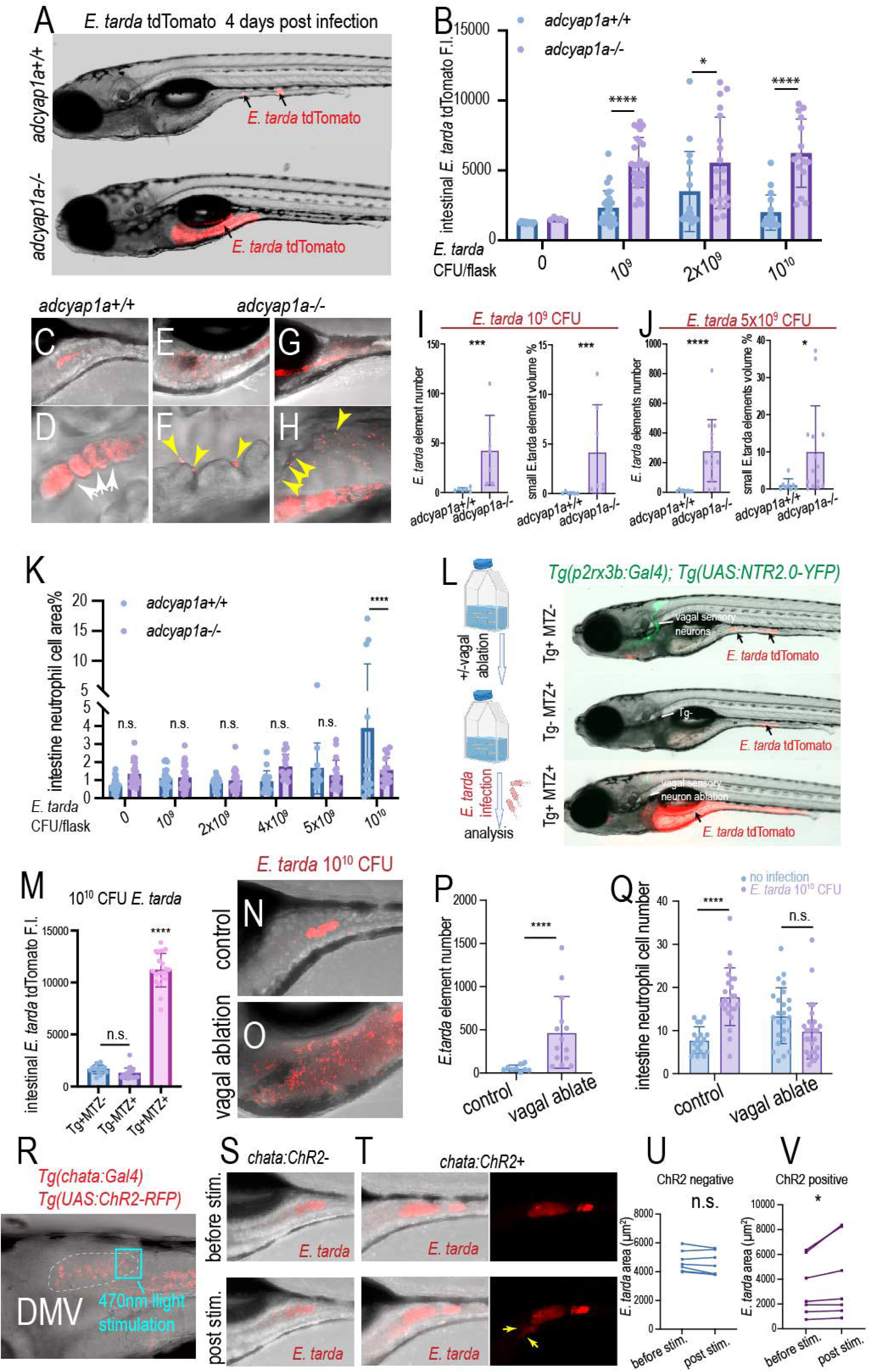
Dysfunction of the Trpa1+EEC-vagal pathway induces *E. tarda* diffusion in the intestine and impairs *E. tarda* clearance. **(A)** Representative images of WT and *adcyap1a*-/- zebrafish that were infected with 10^9^ CFU *E. tarda* tdTomato from 3 dpf to 7 dpf. Note that in WT zebrafish, *E. tarda* forms aggregates, while in adcyap1a-/- zebrafish, *E. tarda* diffuses in the proximal intestine. **(B)** Quantification of intestinal *E. tarda* tdTomato fluorescence in WT and *acyap1a*-/- zebrafish that were infected with different doses of *E. tarda* from 3 dpf to 7 dpf. **(C-D)** Representative image of *E. tarda* in WT zebrafish intestines infected with 10^9^ CFU *E. tarda* from 3 dpf to 7 dpf. C: Z-stack projection. D: Zoomed single-plane image. Note that *E. tarda* forms large aggregates that do not directly touch the intestinal epithelium. **(E-H)** Representative image of *E. tarda* in *adcyap1a*-/- zebrafish intestines infected with 10^9^ CFU *E. tarda* from 3 dpf to 7 dpf. E, G: Z-stack projection. F, H: Zoomed single-plane image. Note that *E. tarda* forms very small clusters or is dispersed as a solitary cell in *adcyap1a*-/-. Yellow arrows in F and H: *E. tarda* that directly touch the intestinal epithelium. **(I-J)** Quantification of *E. tarda* element number and small *E. tarda* element (*E. tarda* aggregates smaller than 300μm^3^) volume ratio in WT and *adcyap1a*-/- zebrafish that are infected with 10^9^ or 5×10^9^ CFU *E. tarda*. **(K)** Quantification of intestinal neutrophil cell area in WT and adcyap1a-/- zebrafish that are infected with different doses of *E. tarda*. **(L)** Epifluorescence images of control and vagal sensory neuron-ablated zebrafish infected with 10^10^ CFU *E. tarda* from 5 dpf to 8 dpf. Vagal sensory neuron ablation was performed via MTZ treatment in *Tg(p2rx3b:Gal4); Tg(UAS:NTR2.0-YFP)* zebrafish from 4 dpf to 5 dpf. Transgenic zebrafish that were not treated with MTZ or non-transgenic zebrafish that were treated with MTZ were used as controls. **(N-O)** Representative confocal z-stack projections of control (transgenic negative zebrafish with MTZ treatment) and vagal sensory neuron-ablated zebrafish that were infected with 10^10^ CFU *E. tarda* from 5 dpf to 8 dpf. **(P)** Quantification of *E. tarda* element number in control and vagal sensory neuron ablated zebrafish. **(Q)** Quantification of intestine neutrophil cell number in WT and adcyap1a-/- zebrafish without or with 10^10^ CFU *E. tarda* colonization. **(R-T)** Stimulating cDMV vagal motor neurons induces *E. tarda* diffusion. **(R)** Confocal image of *Tg(chata:Gal4); Tg(UAS:ChR2-RFP)* zebrafish hindbrain. White box: DMV neurons. Blue box: cDMV region that received 470nm light stimulation. **(S)** Confocal image of *E. tarda* in transgenic negative control zebrafish before and after 470nm light stimulation. **(T)** Confocal image of *E. tarda* in *Tg(chata:Gal4); Tg(UAS:ChR2- RFP)* zebrafish before and after 470nm light stimulation. Note that after stimulating cDMV neurons, some of the E. tarda were dispersed (yellow arrows) from the aggregates. **(U-V)** Quantification of intestine *E. tarda* area before and after 470nm light stimulation in ChR2 transgene-negative or positive zebrafish. B, K, Q: Two-Way ANOVA with Tukey’s post hoc test. (M) One-Way ANOVA with Tukey’s post hoc test. I, J, P: Unpaired Wilcoxon test. U, V: Paired Wilcoxon test. * P<0.05. *** P<0.001. **** P<0.0001.

Larval zebrafish do not have an adaptive immune system, and the immune response to pathogen invasion is primarily carried out by innate immune cells such as neutrophils^46^. As such, we stained uninfected and different doses of *E. tarda* infected zebrafish with myeloperoxidase (MPX), which labels neutrophils. Our results showed that high-dose *E. tarda* colonization significantly enhances neutrophil recruitment to the intestine (**Fig. 8K, Fig. S11**). However, *E. tarda* induced neutrophil recruitment is blunted in *adcyap1a*-/- zebrafish (**Fig. 8K, Fig. S11**). Moreover, our data showed that loss of *adcyap1a* impairs zebrafish survival in response to high-dose *E. tarda* colonization (**Fig. S11**). Together, these data showed that Trpa1+EEC Pacap signaling is required for neutrophil-mediated immune response toward *E. tarda* colonization and enhances survival.

Because Trpa1+EEC Pacap signaling is critical for mediating Trpa1+EEC-vagal communication, we next investigated the role of vagal sensory neurons in the *E. tarda* defense response. To this end, we ablated vagal sensory neurons using the *Tg(p2rx3b:Gal4); Tg(UAS:NTR2.0-YFP)* model above. After vagal sensory neuron ablation, we infected zebrafish with *E. tarda* tdTomato. Like *adcyap1a*-/- zebrafish, we observed that loss of vagal sensory neurons induces *E. tarda* to diffuse in the intestinal lumen (**Fig. 8L, N-P**), and we observed a striking increase in *E. tarda* tdTomato fluorescence in vagal sensory neuron-ablated zebrafish (**Fig. 8L-M**). Using a *Tg(lyz:EGFP)* transgenic line that labels neutrophils, our data also showed that *E. tarda* colonization induced neutrophil recruitment is impaired in vagal sensory neuron ablated zebrafish (**Fig. 8Q**).

Stimulating Trpa1+EECs inhibits a subset of cDMV neurons as shown by our data above. Next, we investigated whether counteracting Trpa1+EEC-induced cDMV neuron inhibition affects *E. tarda* aggregation. To do this, we used a *Tg(chata:Gal4); Tg(UAS:ChR2-RFP)* transgenic zebrafish to express the optogenetic protein Channelrhodopsin-2 (ChR2) in vagal motor neurons. We treated the zebrafish with *E. tarda* tdTomato for 4 days. Following *E. tarda* colonization, 470nm light was delivered to the cDMV region to activate the cDMV neurons, and intestinal *E. tarda* was imaged before and after cDMV stimulation. Our data revealed that in control zebrafish, 470nm light stimulation does not affect E. tarda aggregates (**Fig. 8S, U**). However, when cDMV Tg+ neurons were stimulated, some *E. tarda* dispersed from the aggregates, resulting in an increased *E. tarda* area (**Fig. 8T, V**), suggesting that Trpa1+EEC-induced cDMV neuron inhibition promotes *E. tarda* aggregation.

Collectively, our results demonstrated that the Trpa1+EEC-vagal pathway is important for host defense. Dysfunction of the Trpa1+EEC-vagal pathway induces *E. tarda* diffusion, impairs *E. tarda* clearance, and hinders intestinal neutrophil recruitment.

## Discussion

In this study, using in vivo gut-brain imaging and genetic manipulation in zebrafish, we discovered a distinct non-nutrient sensing Trpa1+EEC-vagal sensory pathway that detects and transmits microbial irritant signals from the intestinal lumen to the brain. Stimulating the Trpa1+EEC-vagal pathway modulates vagal motor neuron activity and increases liver gluconeogenesis. Dysfunction of this gut-brain microbial sensing pathway impairs host immune response and enteric pathogen clearance. Together, our study provides a novel gut-brain neuronal mechanism that is critical for enteric pathogen detection and defense responses.

### The role of the Trpa1+EEC-vagal pathway in sensing microbial irritants

As frontline sensory cells in the intestine, EECs are well known for their nutrient sensory ability. Many studies have shown that EECs express nutrient sensors and can be activated by nutrients such as glucose and fatty acids. In response to nutrient stimulation, EECs secrete hormone peptides and neurotransmitters^14,15^. As such, many EEC-secreted hormones, such as CCK, GLP-1, and PYY, play critical roles in regulating ingestive behavior and metabolism^14,15^. Many of the EEC hormones activate vagal sensory neurons ^14,15^. Communication between EECs and vagal sensory neurons is crucial for controlling food preferences and regulating feeding behavior^25,26,47^.

Compared with EECs’ nutrient-sensing, our understanding of how EECs respond to gut microbial signals remains limited. EECs are heterogeneous, and different EEC subtypes express distinct surface receptors and hormone profiles. Previous studies in mice showed that isovalerate, a bacterial branched-chain fatty acid metabolite, can activate olfactory receptor 558 to stimulate serotonin release from enterochromaffin cells, which then excites the spinal pelvic nerve and exacerbates visceral pain^17,18^. It is well known that short-chain fatty acids from bacterial fiber fermentation can stimulate the short-chain fatty acid receptor FFAR3/4 in GLP-1-expressing EECs to modulate glucose homeostasis and satiation^48^. Similarly, a recent study demonstrated that bacterial flagellin activates PYY-expressing EECs, which then signal to vagal sensory neurons to reduce food intake^16^. These data suggest that certain types of EECs may serve as a novel front-line sensor to perceive and transmit intestinal luminal microbial information. These microbial sensing EECs can then signal to the spinal or vagal sensory neurons to modulate host physiology and behavior.

Like mammals, zebrafish intestinal lumen also harbors complex gut microbiota. Using in vivo EEC calcium imaging, our previous study showed that an opportunistic fish pathogen, *E. tarda*, can activate a subset of Trpa1-expressing EECs^19^. These Trpa1+EECs express Serotonin, suggesting similarities with mammalian enterochromaffin cells. These Trpa1+EECs are not activated by zebrafish commensal bacteria^19^ or nutrients. Stimulating zebrafish with nutrients exclusively activates a subgroup of EECs that do not express Trpa1. Our data suggest that at the EEC level, a segregation is likely present, with certain EEC groups sensing nutrients that are beneficial to the host while other EEC groups, such as Trpa1+EECs, sense chemical or microbial irritants that are harmful. We found that such segregation is also present at the level of vagal sensory neurons and vagal motor neurons. Our study suggests that, like mammals, the zebrafish gut-brain neuronal pathways are meticulously organized and are tuned to specific types of information. Our data demonstrate that specialized EEC and vagal sensory neuron subtypes are utilized to sense and process different enteric information. Certain pathogenic bacterial information in the intestinal lumen can be perceived by a non-nutrient sensing Trpa1+EEC-vagal sensory pathway. Trpa1 is a highly conserved nociceptive receptor that many organisms use to sense harmful environmental stimuli^49^. To survive, organisms need to differentiate and respond differently to harmful information, such as enteric pathogens, than to beneficial information, such as nutrients that are important for life-sustaining and growth. Our study suggests that these two information perceptions are likely segregated along the gut-brain pathway. Such segregation may enable the brain to accurately delegate specialized responses to life-threatening threats when needed.

### Molecular mechanisms underlying Trpa1+EEC-vagal communication

EECs are known to signal to the vagal sensory nerve fibers via paracrine or synaptic mechanisms^15,23^. The EEC-vagal communication is well-studied in the context of nutrient sensing. For example, upon glucose or fatty acid stimulation, CCK-expressing EECs can secrete CCK, which stimulates CCK receptor in vagal sensory neurons. The activated vagal sensory neurons relay signals to the brain to promote satiation and food preference^23,25,26^. Though much is known about EEC and vagal communication in response to nutrients, our understanding of how EECs communicate with vagal sensory neurons in response to microbial stimulants remains very limited.

Our data suggest that harmful irritant bacterial information and beneficial nutrient information are perceived by different EEC subtypes. Interestingly, the Trpa1+EECs that sense microbial irritants and EECs that sense nutrients exhibit different hormone profiles, suggesting that different EEC subtypes may use different packaged signaling molecules to communicate different information to the vagal sensory neurons. Trpa1+EECs do not express CCK, PYY, or GLP-1 hormones that are found in colonial nutrient sensing EECs. Instead, the Trpa1+EECs highly and specifically express Serotonin and Pacap. Our data suggest that Serotonin and Pacap may serve as distinct alarm signals used by Trpa1+EECs to signal the nervous system about the dangers in the intestine.

The role of Serotonin in mediating EEC and vagal sensory neuron communication has been shown in previous studies^50,51^. However, whether other signaling molecules function together with Serotonin to mediate EEC-vagal communication remains unknown. Our study showed that in addition to Serotonin, Pacap is also important to mediate Trpa1+EEC-vagal communication. Pacap is a neuronal peptide broadly expressed in the central and peripheral nervous system. In the central nervous system, Pacap is involved in anxiety and pain response^52,53^. Beyond the nervous system, previous studies showed that Pacap is expressed in mammalian and non-mammalian EECs^54–56^. However, the function of EEC Pacap signaling remains largely unknown. Here, we show that vagal sensory neurons express both HTR3 and Pacap receptor. Trpa1+EEC Pacap synergizes with Serotonin to activate vagal sensory neurons. Loss of Pacap signaling in Trpa1+EECs significantly reduced the number of activated vagal sensory neurons and shortened the vagal sensory neuron calcium elevation duration in response to Trpa1+EEC stimulation. Our study highlights that, in addition to serotonin, PACAP is also critical for mediating Trpa1+EEC-vagal communication. This suggests that a simple one-signaling-molecule, one-receptor model may not be sufficient to recapitulate EEC-vagal communication in vivo. A specific EEC subtype may use multiple signaling molecules to modulate the duration and strength of its communication with vagal sensory neurons.

### The role of the Trpa1+EEC-vagal pathway in defense response

To maintain gut microbial homeostasis, the host evolved diverse mechanisms to monitor gut microbial species in the intestinal lumen and respond accordingly. Much is known about the immune system’s role in mediating gut microbial homeostasis^1^, but our understanding of how other systems sense and respond to gut microbial information remains very limited. The intestine is innervated by the vagal sensory and motor nerve fibers, forming the bidirectional neuronal pathways that mediate gut-brain communication. Vagal sensory neurons are heavily involved in sensing ingested nutrient information, and the vagal innervation is critical to regulating ingestive behavior and nutrient processing^57^. In addition to nutrients, recent studies have demonstrated that vagal sensory neurons can also respond to microbial information. For example, a recent study in a lung infection model demonstrated that vagal sensory neurons could detect Lipopolysaccharides (LPS) from pathogenic bacterial invasion via Toll-like receptor 4 (TLR4)^58^. The activated vagal sensory neurons then transmitted signals to the hypothalamus, which caused severe sickness, behavioral alterations, inflammation, and hypothermia^58^. Interestingly, in another model of systemic LPS injection, it was shown that vagal sensory neurons do not directly detect LPS but are indirectly activated by cytokines released from immune cells in response to LPS injection^8^. The cytokine-induced vagal sensory activation then signals to the NTS and inhibits inflammation^8^. These data suggest that vagal sensory neurons may exhibit pro- and anti-inflammatory effects. These divergent effects may be mediated by different vagal sensory neuron subtypes^59^. Collectively, these recent discoveries established a novel role of vagal sensory neurons as integrators of microbial information^59^. However, it remains unclear how vagal sensory neurons respond to enteric pathogenic bacteria and how vagal microbial sensing affects host defense and gut bacterial physiology.

Our studies showed that vagal sensory neurons can be activated by the pathogenic bacterium *E. tarda* via intestinal epithelium Trpa1+EECs^19^. Here, we further demonstrated that activation of the Trpa1+EEC-vagal sensory pathway is critical for *E. tarda* clearance and host defense. Dysfunction of the Trpa1+EEC-vagal sensory pathway induces *E. tarda* diffusion in the intestinal lumen and impairs intestinal neutrophil recruitment. Trpa1+EEC-vagal sensory neuron activation may promote host defense partially by elevating liver gluconeogenesis and increasing systemic glucose availability, as inhibiting glucose metabolism induces *E. tarda* diffusion and hinders *E. tarda* clearance. Pathogen infection often acutely triggers blood glucose elevation and enhances glucose utilization rate^60,61^. Although chronic hyperglycemia may worsen bacterial infection outcomes^62^, acute blood glucose elevation can benefit host defense by providing energy for tissues to fight pathogen invasion^63^. For example, in response to infection, neutrophils significantly increase glucose uptake and shift from relying heavily on intracellular glycogen at rest to primarily using extracellular glucose^64^. Blocking glucose uptake or glucose metabolism impairs neutrophil chemotaxis, reduces neutrophil reactive oxygen species production, and weakens bacterial killing^65–69^. It was thought that the bacterial infection-induced hyperglycemia is mediated by stress hormones such as epinephrine^70^. However, adrenergic blockade did not prevent bacterial infection induced glucose elevation and glucose metabolism enhancement^71^. Therefore, it is likely that some other mechanisms may contribute to the hyperglycemia response following pathogenic bacterial invasion. Our study suggests that Trpa1+EECs can sense enteric luminal bacterial irritants (live *E. tarda* or their indole metabolites) and promote glucose production via vagal sensory neurons, underscoring the critical roles of gut-brain neuronal mechanisms in metabolic adaptation in response to enteric pathogen invasion.

The vagal motor neurons innervate most of the visceral organs. Afferent vagal sensory neurons and efferent vagal motor neurons form the vago-vagal circuit that is critical for brain control of physiological function. How gut microbiota regulate vagal motor neuron activity remains largely unknown. Vagal motor neuron activation dampens systemic inflammation via the vagal-spleen axis or directly suppresses intestinal immune cells via vagal motor neuron acetylcholine signaling^72^. Our data showed that *E. tarda* or *E. tarda* indole potently inhibits zebrafish vagal motor neurons that innervate visceral organs. *E. tarda* or *E. tarda* indole induced vagal motor neuron inhibition may be critical to lift vagal-mediated anti-inflammatory effects and prepare the organism to respond appropriately to pathogens in the intestinal lumen. Vagal sensory neurons do not directly signal to vagal motor neurons in the DMV, but indirectly modulate DMV activity through neurons in the NTS^73^. Motor neurons in the DMV are tonically and phasically inhibited by gamma-aminobutyric acid (GABA) releasing neurons in the NTS^74,75^. In addition to GABAergic inhibition, vagal motor neurons in the DMV can also be inhibited by other neuronal peptides such as Galanin^76^. How Trpa1+EEC-vagal sensory activation inhibits vagal motor neuron activity in zebrafish is an intriguing question that requires future studies.

One striking phenotype revealed by our studies is that Trpa1+EECs and vagal sensory neurons not only reduce *E. tarda* burden but also alter *E. tarda* physiology. In wild-type zebrafish, *E. tarda* forms large aggregates and is not motile. These large E. tarda aggregates reside in the lumen and do not touch the intestinal epithelium. Without Trpa1+EEC Pacap signaling or vagal sensory neurons, *E. tarda* is dispersed as single bacterial cells or as very small clusters that are diffuse in the intestinal lumen. These dispersed *E. tarda* cells or small *E. tarda* clusters are motile and directly contact the intestinal epithelium. These findings suggest that *E. tarda* exhibit different physiological phenotypes in the intestine of WT zebrafish or zebrafish with dysfunctional Trpa1+EEC-vagal signaling. The large *E. tarda* aggregates in WT zebrafish may represent biofilm-like clumps, structures bacteria use to protect themselves from environmental stress or to invade immune systems^77,78^. Our data showed that in response to *E. tarda* invasion, increased immune response and neutrophil infiltration were present in WT zebrafish but not in zebrafish with dysfunctional Trpa1+EEC-vagal signaling. The increased immune response in the WT zebrafish intestine may induce *E. tarda* to form large biofilm-like aggregates. In addition to the immune system, activation of the Trpa1+EEC-vagal sensory pathway may also engage other host defense systems to modify *E. tarda* physiology. Our previous study showed that Trpa1+EECs also signal to the enteric neurons and promote intestinal movement^19^. Whether and how the enteric nervous system is involved in the Trpa1+EEC-vagal mediated host defense is an intriguing question that can be studied in the future.

Together, our study suggests that the Trpa1+EEC-vagal sensory pathway represents a novel mechanism by which the brain monitors certain irritant microbial information in the intestinal lumen. Stimulating the Trpa1+EEC-vagal pathway transmits signals to the brain to coordinate physiological and immune changes to fight pathogenic bacterial invasion. The Trpa1+EEC-vagal pathway may enable early detection of the enteric pathogen before the pathogen crosses the intestinal barrier and spreads systemically. Thus, this pathway may benefit the host during acute enteric pathogen invasion, and dysfunction of the Trpa1+EEC-vagal signaling impairs host defense and pathogen clearance. However, chronic activation of this pathway may impair vagal motor neuron function and disrupt vagal motor control of visceral organ function and physiological homeostasis.

## Methods

### Zebrafish strains and husbandry

All zebrafish experiments conformed to the US Public Health Service Policy on Humane Care and Use of Laboratory Animals, using protocol number 2021A00000091A1 approved by the Institutional Animal Care and Use Committee of the Ohio State University. Conventionally-reared adult zebrafish were reared and maintained on a recirculating aquaculture system using established methods ^79^. For experiments involving conventionally-raised zebrafish larvae, adults were bred naturally in system water and fertilized eggs were transferred to 100mm petri dishes containing ∼25mL of egg water at approximately 6 hours post-fertilization. The resulting larvae were raised under a 14h light/10h dark cycle in an air incubator at 28°C at a density of 2 larvae/ml water. All the experiments performed in this study ended at 7 dpf unless specifically indicated. The zebrafish lines used in this study are listed in **Table 1**. All lines were maintained on a EKW background.

To generate *adcyap1a* mutant zebrafish, we used CRISPR/Cas9 technique as described previously^19^. In brief, a guide RNA that targets 5’-GAACTAGGAGCCCATAGATG-3’ in exon 1 of *adcyap1a* gene shortly after the ATG start site was used. crRNA and TracrRNA were directly ordered from IDT. The gRNA was co-injected with Cas9-NLS protein (PNA-Bio PC01) into zebrafish eggs at one cell stage. CRISPR efficiency in F0 zebrafish was assessed via heteroduplex mobility shift assay. A +5 stable mutant and its genomic sequence (**Fig. S7**) were identified in F2. The insertion created a new HaeIII restriction enzyme cutting site. Genotyping of adcyap1a-/- zebrafish can be performed via PCR using forward primer 5’-GGGAATCGTACTTCATTTATAGCAA-3’ and reverse primer 5’-TTCCTCAGTAAAATGTGCTCAAAG-3’. The resulting PCR product is digested with the HaeIII enzyme (NEB R0108S) and analyzed via DNA electrophoresis. WT zebrafish present a single 405bp band, and adcyap1a-/- zebrafish have two bands.

### Bacterial culture and genetic manipulation

*E. tarda* FL6-60 strain was cultured using Tryptic Soy Agar (TSA) plates or Tryptic Soy Broth (TSB) media at 30 °C. The Tn7 transposase insertion approach was used to insert the tdTomato gene at the attTn7 site into the bacterial glms gene site, as reported previously^44^. The pTNS2 plasmid (Addgene 64968) and pTn7-TS-tdTomato plasmid (Addgene 117391) were ordered from Addgene. pTNS2 and pTn7 plasmids were then transformed into *E. coli* sm10λ pir. The transformed *E. coli* sm10λ pir were then mated with *E. tarda* FL6-60. Antibiotics and temperature were then used select *E. tarda* FL6-60 that carries tdTomato insertion. 16S sequencing and PCR based genomic sequencing were used to confirm the resulting *E. tarda* FL6-60 strain.

### Gnotobiotic zebrafish manipulation and *E. tarda* colonization

To determine host response to *E. tarda*, the conventionally raised zebrafish were stimulated with different doses of *E. tarda*. *Edwardsiella tarda* FL6-60 with tdTomato integrated into the chromosome was struck out onto a TSA + Gentamycin agar plate and incubated at 30°C overnight. A single colony was used to inoculate 5mL of TSB, which was then grown at 30°C shaking overnight. This was used to inoculate 40mL of TSB at a 1:1000 dilution. This culture was grown at 30°C shaking to a concentration of ∼0.500-0.600 OD600 determined by a V-1200 Spectrophotometer (VWR). *E. tarda* OD600 and *E. tarda* CFU lineage relationship was measured, and the approximate *E. tarda* CFU/mL of this culture was calculated accordingly. This culture was spun down at 3220xg for 10-15 minutes to pellet the cells and was then washed 3 times with zebrafish egg water. After the final wash, the cells were resuspended in 10mL egg water, which was then used to inoculate groups of 30 3dpf zebrafish kept in 25mL of egg water in T25 cell culture flasks at the desired *E. tarda* concentration. These fish were kept at 28°C, with 50% water changes every 24 hours, making note of and removing any deceased fish.

To determine how commensal bacteria and *E. tarda* affect zebrafish liver gluconeogenesis, we used a zebrafish gnotobiotic approach. Germ-free (GF) zebrafish were generated as previously described^21^. Germ-free zebrafish eggs were maintained in cell culture flasks with sterile gnotobiotic zebrafish media (GZM) at a density of 1 larvae/ml. From 3 dpf to 7 dpf, a 60% daily media change, and a feeding of 250µl of 1% newborn fish food (Ultra Fresh Ltd.) feeding were performed as described. To colonize GF zebrafish with commensal microbiota, 15 mL filtered system water (5μm filter, SLSV025LS, Millipore, final concentration of system water ∼30%) was inoculated into flasks containing germ-free zebrafish in GZM at 3 dpf, when the zebrafish normally hatch from their protective chorions. The same feeding and media change protocol was followed as for germ free zebrafish. Microbial colonization density was determined via Colony Forming Unit (CFU) analysis. To colonize GF zebrafish with *E. tarda*, 250 μl overnight *E. tarda* culture was harvested, washed three times with sterile GZM, and then inoculated into flasks containing 3 dpf germ free zebrafish. The same feeding and media change protocol was followed as for germ free zebrafish. Microbial colonization density was determined via Colony Forming Unit (CFU) analysis.

### In vivo enteroendocrine cell calcium imaging

All the in vivo imaging was performed using the Nikon AXR confocal. When imaging the EEC cellular and mitochondrial calcium activity using the *Tg(neurod1:Gcamp6f); Tg(neurod1:mitoRGECO)* zebrafish, the zebrafish were not anesthetized due to the effects of Tricaine in activating EECs. 7 dpf zebrafish larvae were mounted in the 35mm confocal dish using 1% low melting agar. In the experiments where chemical stimulants were applied a small window was cut in front of the zebrafish, which allowed the zebrafish’s mouth to be exposed. 1ml of chemical stimulants was applied to the zebrafish as described previously. The zebrafish intestine was imaged before and after the chemical stimulants. The following chemical stimulants were used: 1.6mM linoleic acid, 10% glucose, and 1mM indole. The image was captured using a 20× LWD water objective lens and the resonant scanner.

For the image analysis to assess the EEC calcium activity, the images were first aligned using the Nikon NLS element software 3D segmentation and 3D tracking function. The EECs are selected and segmented by the Gcamp fluorescence based on the threshold intensity. The threshold was applied to identify more than 95% of EECs, and the threshold was consistent across one experiment. Following threshold identification, image smoothing was applied with parameter 0.85µm, objects less than 1µm were removed, and the object with a distance of more than 0.22µm was separated. Following segmentation, for each object unit, the mean Gcamp6f fluorescence intensity, mean mitoRGECO fluorescence intensity and object volume were measured. The objects with a volume outside of 100 µm^3^-1000 µm^3^ were filtered out. Following segmentation, the individual object units in different time frames were traced and tracked via the NLS element 3D-object tracing software. Due to the issues of gut motility, not every EEC in the zebrafish can be successfully traced throughout the time course. The mean fluorescence intensity of the individual EEC in each time frame will be calculated. The cluster 3.0 software was used to perform the clustering analysis of the EECs that exhibit different temporal calcium dynamics.

### Simultaneous optic stimulation and brain calcium imaging

To perform simultaneous optic stimulation in the zebrafish and brain calcium imaging, a digital micromirror device (DMD) was installed in the AXR^80^. DMD is a semiconductor-based micro-electromechanical system (MEMS) containing thousands of of tiny, individually tilting aluminum mirrors. When a mirror tilts to the “on” position, it reflects light through a projection lens to stimulate a pixel of the selected ROI. The DMD device was coupled with an optical light source that can deliver selective wavelengths of fluorescence light (405nm, 470nm, and 590nm). Note that the stimulation light source and the confocal laser light source are separate, which allows simultaneous light stimulation and confocal imaging. 7 dpf zebrafish larvae were mounted in 1% low melting agar. Zebrafish were not anesthetized as anesthetics would significantly affect neuronal calcium. A region of interest will be defined, and the light stimulation protocol will be set up using the Nikon NLS element software. To stimulate Trpa1+EECs in the intestine, zebrafish were pretreated with 10μM Optovin (Cayman Chemical, catalog number 14851) for 2 hours. Following optovin treatment, zebrafish were mounted on their side and positioned in the AXR confocal. To stimulate caudal DMV neurons, *Tg(chata:Gal4); Tg(UAS:ChR2-RFP)* zebrafish were used. A 20× water objective lens was used. 405nm light was selectively delivered to the middle intestine (Fig. S3). The vagal calcium activity was recorded before, during, and after light stimulation. *Tg(elavl3:H2B-Gcamp6s)* was be used to record vagal sensory neuron calcium activity. *Tg(chata:Gal4); Tg(UAS:Gcamp6s)* was be used to record vagal motor neuron calcium activity.

Following the calcium recording, the images were denoised and aligned using the NLS Element software. Neurons were manually segmented and the temporal Gcamp fluorescence for individual segmented neurons was measured. The Gcamp fluorescence was normalized to T0.

### Measurement of liver gluconeogenesis and glucose production

Zebrafish liver gluconeogenesis is measured via the *Tg(pck1:venus; cryaa:RFP)* reporter line. Following treatment, live zebrafish were mount in 3% methylcellulose. The pck1:venus fluorescence is imaged using the Zeiss Axio Zoom V16 upright epifluorescence microscope via the EGFP filter. To measure zebrafish glucose, we used a previously published method^81^. In brief, zebrafish were pooled and lysed using the tissue tearer, and the supernatant was collected. Glucose concentration in the supernatant was measured via the colorimetric assay using the glucose assay kit (Abcam ab65333).

### Immunofluorescence staining and chemical treatment

Whole mount immunofluorescence staining was performed as previously described^24^. In brief, ice cold 2.5% formalin was used to fix zebrafish larvae overnight at 4°C. The samples were then washed with PT solution (PBS+3%Triton-100). The skin and remaining yolk were then removed using forceps under a dissecting microscope. The deyolked samples were then permeabilized with methanol for more than 2 hours at -20°C. Samples were then blocked with 4% BSA at room temperature for more than 1 hr. The primary antibody was diluted in PT solution and incubated at 4°C for more than 24 hrs. Following primary antibody incubation, the samples were washed with PT solution and incubated overnight with secondary antibody with Hoechst 33342 for DNA staining. Imaging was performed with Nikon AXR confocal using the 20× of 40× water immersion lens. The primary antibodies that were used are listed in **Table 1**. The secondary antibodies in this study were from Alexa Fluor Invitrogen and were used at a dilution of 1:250.

The chemicals used in this manuscript are included in **Table 1**. In general, the chemicals were dissolved in a DMSO stock solution and diluted to the desired concentration right before use. DMSO was used as a control. Tropisetron (200 μM), P8 (50 μM) were used in the experiments. For indole stimulation, 1 mM was used for acute stimulation to examine the effects on EEC and vagal activity. 100μM were used for prolonged stimulation. For all the other tryptophan metabolites that were tested in Fig. 6, 100μM was used.

### HCR In Situ Hybridization

Hybridization Chain Reaction-Fluorescence In Situ Hybridization (HCR-FISH) was performed by using the HCRTM RNA-FISH kit and adapting its standard protocol (Molecular Instruments, United States). adcyap1r1a and htr3a probes were customized ordered from the same vendor. Briefly, 7 dpf healthy larvae of WT EK fish were fixed with cold fixing buffer (2.5% formaldehyde, 0.25% TritonX-100 in PEM buffer) for 48 hours in 4°C. The dehydration, permeabilization, detection, and signal amplification steps were then performed according to kit protocol, with a modification on proteinase K digestion of using 3ug/ml proteinase K digesting for 20 minutes instead of the suggested step in the protocol. After the final step of amplification, incubate sample in a DNA staining solution (1% BSA, 1% DMSO, 0.3% TritonX-100, 1xPBS) with 1:400 of Hoechst 33342 (Thermo Fisher Scientific, United States) overnight in 4°C. DNA stain was removed the next day and the samples were washed with 1xPBS + 0.3% TritonX-100 for three times for five minutes each. The stained samples were stored in 1xPBS + 0.02% sodium azide at 4°C until ready to image.

### Statistical analysis

Preliminary experiments were performed to evaluate variance and effects. The data points in most figures represent individual zebrafish unless specified otherwise. For most experiments comparing results between 2 groups, the unpaired Student’s t-test will be used, and significance will be assigned to P values <0.05 with two-tailed calculation. For experiments in Figure 8 that are involved in *E. tarda* element quantification, unpaired or paired Wilcoxon tests were used. When comparing results among 3 or more groups, ANOVA will be used, followed by the Tukey-Kramer or Sidak’s multiple-comparison posttest, and significance will be assigned at P <0.05.

## Supplemental Figure legends

**Figure S1. Trpa1+EECs are separate from the other EEC subgroups. (A-G)** UMAP plots of the zebrafish intestine single-cell RNA sequencing dataset generated by Wen J. et al., 2021^20^ showing the expression of different genes. *neurod1*: transcription factor that labels all EECs. *ccka* (*cholecystokinin a*)*, gcga* (*preproglucagon a*)*, pyyb* (*peptide YY b*): Hormones that are expressed in nutrient-sensing EECs. Note that Trpa1+EECs do not express these hormones. *Sglt1* (*sodium-glucose co-transporter 1*) and *ffar1* (*free fatty acid receptor 1*): nutrient sensors that sense glucose and long-chain/medium-chain fatty acids.

**Figure S2. Analysis of zebrafish EEC activity in vivo.** (A) Schematic model showing the experimental setup and EEC calcium imaging in response to nutrient stimulants. (B) Time-lapse image frame showing EEC calcium response after linoleic acid stimulation. (C) Quantification of four EEC calcium profiles after linoleic acid stimulation. Top two: linoleic acid-activated EECs. Bottom two: the EECs that are not activated by linoleic acid. (D) Heatmap and clustering analysis showing EEC calcium response.

**Figure S3. Regional specificity of light stimulation.** *Tg(neurod1:mitoEOS)* zebrafish line was used. In this model, mitoEOS was expressed in EECs, pancreatic islet cells, and brain neuronal cells. mitoEOS is a green fluorescent protein. 405nm light can convert mitoEOS from green to red. We delivered the 405nm light specifically to the yellow-boxed region in the zebrafish intestine. Note that only EECs inside the yellow-boxed region were converted to red, while other cells remained green.

**Figure S4. Stimulating Trpa1+EECs activates a subset of vagal sensory neurons. (A)** Experimental setup. *Tg(neurod1:Gcamp6s)* zebrafish were pretreated with Optovin. 405nm light was delivered to the intestine to activate Trpa1+EECs, and vagal sensory neuron calcium activity was recorded simultaneously. **(B-E)** Time-lapse image frames showing vagal sensory neuron calcium activity before and after Trpa1+EEC stimulation. Yellow circles: three vagal sensory neurons that increased Ca^2+^ activity following Trpa1+EEC stimulation. **(F)** Quantification of the three circled vagal sensory neurons’ calcium profiles before and after Trpa1+EEC stimulation. **(G)** Heatmap and clustering analysis showing vagal sensory neuron calcium response before and after Trpa1+EEC stimulation. Vagal sensory neuron calcium activity was measured by *neurod1:Gcamp6s*. **(H-J)** 405nm light stimulation in control zebrafish that are not treated with Optovin does not activate vagal sensory neurons. **(I)** Time-lapse image frames showing vagal sensory neuron calcium activity before and after 405nm light stimulation in zebrafish that are not treated with Optovin. **(J)** Heatmap and clustering analysis showing vagal sensory neuron calcium response before and after 405nm light stimulation in zebrafish that are not treated with Optovin.

**Figure S5. Glucose activates a subset of zebrafish vagal sensory neurons.** The vagal sensory neuron calcium activity was measured in *Tg(elavl3:H2B-Gcamp6s)*. (A-D) 3D projection of the vagal sensory ganglion before and 5 minutes after glucose or water stimulation. Note that glucose, but not water, activates a subset of vagal sensory neurons. **(E)** Heatmap showing vagal sensory neuron calcium response before and after glucose stimulation.

**Figure S6. HTR3a and adcyap1r1a were expressed in cranial sensory neurons.** The single-cell RNA sequencing dataset was generated by Sur A. et al., 2023^27^. **(A-B)** UMAP plots showing the expression of *htr3a* and *adcyap1r1a* in zebrafish neuronal cells. Boxes in A and B: cranial sensory neurons. Note that htr3a is highly and specifically expressed in cranial sensory neurons but not in other neuronal cells. Adcyap1r1a is broadly expressed in different neuronal cells. **(C)** Expression of different serotonin receptors in cranial sensory neurons. Note that only htr3a, but not other serotonin receptors, was expressed in the cranial sensory neurons.

**Figure S7. A new *adcyap1a*-/- zebrafish model. (A)** DNA sequence of WT and *adcyap1a*-/- zebrafish generated by CRISPR-Cas9. gRNA was designed to target the highlighted sequence in Exon 1 shortly after the ATG start codon. The mutant zebrafish has a 5-nucleotide insertion, which leads to a frameshift and a premature stop codon. **(B-C)** Immunofluorescence staining of PACAP in WT and *adcyap1a*-/- zebrafish intestine. Note that PACAP is not expressed in EECs but is still present in enteric neurons in adcyap1a-/- zebrafish. **(D-E)** Expression of *adcyap1a* and *adcyap1b* in whole zebrafish larvae cells. Note that adcyap1a and adcyap1b have different expression patterns. Adcyap1a is specifically expressed in EECs, while adcyap1b is broadly expressed in central and peripheral neurons.

**Figure S8. A cluster of vagal motor neurons in the caudal DMV displays high basal activity. (A-A’)** Z-stack projection of vagal sensory and motor neurons. Vagal sensory neurons were labeled by *Tg(p2rx3b:mem-mCherry)*, and vagal motor neuron calcium was indicated by *Tg(chata:Gal4);Tg(UAS:Gcamp6s)*. **(B)** DMV neuron calcium activity. Note that a group of vagal motor neurons in the caudal DMV display higher calcium activity than the remaining DMV neurons. **(C)** Temporal calcium profiles of two vagal motor neurons circled in B. Note that the circled caudal DMV neuron maintained a higher GCaMP fluorescence level during the imaging phase.

**Figure S9. Indole or *E. tarda* decreases caudal DMV neuron activity. (A-B)** Z-stack projection of vagal motor neurons in 7dpf zebrafish that are treated with DMSO or indole for 12 hours. Vagal motor neurons were labeled by *Tg(chata:Gal4); Tg(UAS:Gcamp6s)*. **(C-D)** Quantification of the rostral and caudal DMV neuron Gcamp fluorescence in DMSO and indole treated zebrafish. **(E-H)** Caudal DMV vagal motor neuron activity in *E. tarda* or commensal microbiota colonized zebrafish. Germ-free (GF) zebrafish were colonized with *E. tarda* or commensal microbiota from 3dpf to 7dpf. Neuronal activity was analyzed by pERK staining. *Tg(isl1:EGFP)* was used to label DMV neurons. **(F-G’)** Z-stack projection of GF and *E. tarda* colonized zebrafish DMV. (H) Quantification of caudal DMV neuron activity in GF, *E. tarda* colonized, and commensal microbiota colonized (CV) zebrafish. Student t’sT-test was used in C and D. One-Way ANOVA with Tukey post hoc test was used in H. **** P<0.0001.

**Figure S10. Zebrafish hepatocytes do not express PACAP receptors. (A-F)** UMAP plots of the zebrafish intestine single-cell RNA sequencing dataset generated by Sur A. et al., 2023^27^, showing the expression of *adcyap1r1a* and *adcyap1r1b* in zebrafish cells. (A, D) UMAP plot of all zebrafish cells. (B, E) UMAP plot of zebrafish neuronal cells. (C, F) UMAP plot of zebrafish endoderm cells. **(G)** UMAP plot of fabp10a, a key transcription factor of zebrafish liver in endoderm cells. Note that both *adcyap1r1a* and *adcyap1r1b* are strongly expressed in zebrafish neurons but are not expressed in the liver. **(H-H’’)** HCR-ISH staining showing the expression of *adcyap1r1a* and *htr3a* in 7 dpf zebrafish. Note that *adcyap1r1a* is broadly expressed in the nervous system, including vagal sensory neurons, while htr*3a* is specifically expressed in vagal sensory neurons. **(I-I’’)** Zoomed view showing that neither adcyap1r1a nor htr3a are expressed in the liver.

**Figure S11. Loss of adcyap1a impairs *E. tarda-*induced intestinal neutrophil recruitment and survival. (A)** Distribution of zebrafish with different *E.tarda* phenotypes in the intestine. WT or *adcyap1a*-/- zebrafish were infected with different doses of *E. tarda* from 3 dpf to 7 dpf and were imaged at 7 dpf. Blue: no *E. tarda* presence. Yellow: *E. tarda* as big aggregates. Red: *E. tarda* diffused in the intestinal lumen. **(B)** Survival of WT and adcyap1a-/- zebrafish that were colonized with *E. tarda*. Zebrafish were infected with 10^10^ CFU *E.tarda* from 3 dpf to 6 dpf or from 3 dpf to 7 dpf. 60 WT and 60 adcyap1a-/- zebrafish were analyzed. **(C-D’)** Epifluorescence images of representative WT zebrafish without or with 10^10^ CFU *E. tarda* infection. Neutrophil cells were labeled by MPX staining. **(E)** Zoomed image showed the neutrophils near the red fluorescently labeled *E. tarda*. **(F-G ‘)** Epifluorescence images of representative adcyap1a-/- zebrafish without or with 10^10^ CFU *E. tarda* colonization. **(H)** Zoomed image shows the neutrophils near the red fluorescently labeled *E. tarda*.

## Supplemental videos and table

**Video 1: Light-induced Trpa1+EEC activation.** *Tg(neurod1:Gcamp6f)* was used to image EEC calcium. Zebrafish were treated with Optovin. 405nm light was delivered to the imaged region.

**Video 2: Stimulating Trpa1+EECs activates a subset of vagal sensory neurons.** *Tg(elavl3:H2BGcamp6s)* was used to image vagal sensory neuron calcium. Zebrafish were treated with optovin. 405nm light was delivered to the intestine when the vagal sensory ganglion was imaged.

**Video 3: Glucose activates a subset of zebrafish vagal sensory neurons.** *Tg(elavl3:H2BGcamp6s)* was used to image vagal sensory neuron calcium. A time-lapse z-stack projection was captured. Vagal sensory ganglion calcium was imaged before and after glucose stimulation.

**Video 4: Water does not activate zebrafish vagal sensory neurons.** *Tg(elavl3:H2BGcamp6s)* was used to image vagal sensory neuron calcium. A time-lapse z-stack projection was captured. Vagal sensory ganglion calcium was imaged before and after water stimulation.

**Video 5: Vagal sensory central projection and dorsal motor nucleus of the vagus in the zebrafish hindbrain.** *Tg(p2rx3b:EGFP)* was used to label vagal sensory neuron central projection. *Tg(chata:Gal4); Tg(UAS:NTR-mCherry)* was used to label the vagal motor neurons in the DMV. The zebrafish hindbrain region was imaged.

**Video 6: Vagal motor neuron response to linoleic acid stimulation.** *Tg(chata:Gal4); Tg(UAS:Gcamp6s)* was used to image vagal motor neuron calcium. A time-lapse z-stack projection was captured. Vagal motor neurons in the DMV were imaged before and after linoleic acid stimulation.

**Video 7: Vagal motor neuron response to indole stimulation.** *Tg(chata:Gal4); Tg(UAS:Gcamp6s)* was used to image vagal motor neuron calcium. A time-lapse z-stack projection was captured. Vagal motor neurons in the DMV were imaged before and after indole stimulation.

**Video 8: *E. tarda* aggregates in WT zebrafish intestine.** WT zebrafish was infected with *E. tarda* tdTomato for 4 days. A time-lapse z-stack projection was captured.

**Video 9: *E. tarda* elements in *adcyap1a*-/- zebrafish intestine.** *adcyap1a*-/- zebrafish were infected with *E. tarda* tdTomato for 4 days. A time-lapse z-stack projection was captured.

**Table 1: Zebrafish lines, primary antibodies, and chemicals used in this paper.**

## References

1 Chung, H. & Kasper, D. L. Microbiota-stimulated immune mechanisms to maintain gut homeostasis. Curr Opin Immunol 22, 455–460, doi:10.1016/j.coi.2010.06.008 (2010).

2 Lee, J. Y., Tsolis, R. M. & Baumler, A. J. The microbiome and gut homeostasis. Science 377, eabp9960, doi:10.1126/science.abp9960 (2022).

3 Rogers, A. P., Mileto, S. J. & Lyras, D. Impact of enteric bacterial infections at and beyond the epithelial barrier. Nat Rev Microbiol 21, 260–274, doi:10.1038/s41579-022-00794-x (2023).

4 Blouet, C. & Schwartz, G. J. Brain Nutrient Sensing: A Unifying Framework. Annu Rev Physiol 88, 203–227, doi:10.1146/annurev-physiol-042324-100329 (2026).

5 Gomez-Pinilla, F. Brain foods: the èects of nutrients on brain function. Nat Rev Neurosci 9, 568–578, doi:10.1038/nrn2421 (2008).

6 Koren, T. et al. Insular cortex neurons encode and retrieve specific immune responses. Cell 184, 5902–5915 e5917, doi:10.1016/j.cell.2021.10.013 (2021).

7 Ilanges, A. et al. Brainstem ADCYAP1(+) neurons control multiple aspects of sickness behaviour. Nature **609**, 761–771, doi:10.1038/s41586-022-05161-7 (2022).

8 Jin, H., Li, M., Jeong, E., Castro-Martinez, F. & Zuker, C. S. A body-brain circuit that regulates body inflammatory responses. Nature 630, 695–703, doi:10.1038/s41586-024-07469-y (2024).

9 Sparling, A. C., Florsheim, E. B. & Sullivan, Z. A. Scaling immunity: sickness as a host defense strategy. Trends Immunol 47, 460–462, doi:10.1016/j.it.2026.03.015 (2026).

10 Wang, A. et al. Opposing Èects of Fasting Metabolism on Tissue Tolerance in Bacterial and Viral Inflammation. Cell 166, 1512–1525 e1512, doi:10.1016/j.cell.2016.07.026 (2016).

11 Dantzer, R. Evolutionary Aspects of Infections: Inflammation and Sickness Behaviors. Curr Top Behav Neurosci 61, 1–14, doi:10.1007/7854_2022_363 (2023).

12 Goehler, L. E. et al. Activation in vagal àerents and central autonomic pathways: early responses to intestinal infection with Campylobacter jejuni. Brain Behav Immun **19**, 334–344, doi:10.1016/j.bbi.2004.09.002 (2005).

13 Riley, T. P., Neal-McKinney, J. M., Buelow, D. R., Konkel, M. E. & Simasko, S. M. Capsaicin-sensitive vagal àerent neurons contribute to the detection of pathogenic bacterial colonization in the gut. J Neuroimmunol 257, 36–45, doi:10.1016/j.jneuroim.2013.01.009 (2013).

14 Furness, J. B., Rivera, L. R., Cho, H. J., Bravo, D. M. & Callaghan, B. The gut as a sensory organ. Nat Rev Gastroenterol Hepatol 10, 729–740, doi:10.1038/nrgastro.2013.180 (2013).

15 Gribble, F. M. & Reimann, F. Function and mechanisms of enteroendocrine cells and gut hormones in metabolism. Nat Rev Endocrinol 15, 226–237, doi:10.1038/s41574-019-0168-8 (2019).

16 Liu, W. W. et al. A gut sense for a microbial pattern regulates feeding. Nature 645, 729–736, doi:10.1038/s41586-025-09301-7 (2025).

17 Bellono, N. W. et al. Enterochromàin Cells Are Gut Chemosensors that Couple to Sensory Neural Pathways. Cell 170, 185–198 e116, doi:10.1016/j.cell.2017.05.034 (2017).

18 Bayrer, J. R. et al. Gut enterochromàin cells drive visceral pain and anxiety. Nature 616, 137–142, doi:10.1038/s41586-023-05829-8 (2023).

19 Ye, L. et al. Enteroendocrine cells sense bacterial tryptophan catabolites to activate enteric and vagal neuronal pathways. Cell Host Microbe **29**, 179–196 e179, doi:10.1016/j.chom.2020.11.011 (2021).

20 Wen, J. et al. Fxr signaling and microbial metabolism of bile salts in the zebrafish intestine. Sci Adv 7, doi:10.1126/sciadv.abg1371 (2021).

21 Alsudayri, A. et al. Gut microbiota regulate maturation and mitochondrial function of the nutrient-sensing enteroendocrine cell. Development 151, doi:10.1242/dev.202544 (2024).

22 Ye, L. et al. High fat diet induces microbiota-dependent silencing of enteroendocrine cells. Elife 8, doi:10.7554/eLife.48479 (2019).

23 Kaelberer, M. M. et al. A gut-brain neural circuit for nutrient sensory transduction. Science 361, doi:10.1126/science.aat5236 (2018).

24 Runyon, A. et al. Enteroendocrine cells wire the gut-brain vagal axis. Proc Natl Acad Sci U S A 123, e2600339123, doi:10.1073/pnas.2600339123 (2026).

25 Tan, H. E. et al. The gut-brain axis mediates sugar preference. Nature 580, 511–516, doi:10.1038/s41586-020-2199-7 (2020).

26 Li, M. et al. Gut-brain circuits for fat preference. Nature 610, 722–730, doi:10.1038/s41586-022-05266-z (2022).

27 Sur, A. et al. Single-cell analysis of shared signatures and transcriptional diversity during zebrafish development. Dev Cell 58, 3028–3047 e3012, doi:10.1016/j.devcel.2023.11.001 (2023).

28 Kutz, K. Pharmacology, toxicology and human pharmacokinetics of tropisetron. Ann Oncol 4 **Suppl 3**, 15–18, doi:10.1093/annonc/4.suppl_3.s15 (1993).

29 Takasaki, I. et al. In Silico Screening Identified Novel Small-molecule Antagonists of PAC1 Receptor. J Pharmacol Exp Ther 365, 1–8, doi:10.1124/jpet.117.245415 (2018).

30 Takasaki, I. et al. The novel small-molecule antagonist of PAC1 receptor attenuates formalin-induced inflammatory pain behaviors in mice. J Pharmacol Sci 139, 129–132, doi:10.1016/j.jphs.2018.11.011 (2019).

31 Randlett, O. et al. Whole-brain activity mapping onto a zebrafish brain atlas. Nat Methods 12, 1039–1046, doi:10.1038/nmeth.3581 (2015).

32 Barsh, G. R., Isabella, A. J. & Moens, C. B. Vagus Motor Neuron Topographic Map Determined by Parallel Mechanisms of hox5 Expression and Time of Axon Initiation. Curr Biol 27, 3812–3825 e3813, doi:10.1016/j.cub.2017.11.022 (2017).

33 Ruetten, V. M. S. et al. Imaging cellular activity across all organs reveals body-wide circuits. Nature, doi:10.1038/s41586-026-10979-6 (2026).

34 Isabella, A. J., Barsh, G. R., Stonick, J. A., Dubrulle, J. & Moens, C. B. Retinoic Acid Organizes the Zebrafish Vagus Motor Topographic Map via Spatiotemporal Coordination of Hgf/Met Signaling. Dev Cell 53, 344–357 e345, doi:10.1016/j.devcel.2020.03.017 (2020).

35 Pocai, A., Obici, S., Schwartz, G. J. & Rossetti, L. A brain-liver circuit regulates glucose homeostasis. Cell Metab 1, 53–61, doi:10.1016/j.cmet.2004.11.001 (2005).

36 Brito, C. F. et al. Vagus Nerve Mediated Liver-Brain-Axis Is a Major Regulator of the Metabolic Landscape in the Liver. Int J Mol Sci 26, doi:10.3390/ijms26052166 (2025).

37 Metz, C. N. & Pavlov, V. A. Vagus nerve cholinergic circuitry to the liver and the gastrointestinal tract in the neuroimmune communicatome. Am J Physiol Gastrointest Liver Physiol 315, G651–G658, doi:10.1152/ajpgi.00195.2018 (2018).

38 Kwon, E. et al. Optogenetic stimulation of the liver-projecting melanocortinergic pathway promotes hepatic glucose production. Nat Commun 11, 6295, doi:10.1038/s41467-020-20160-w (2020).

39 Wang, L. et al. Identification of Insulin-Responsive Transcription Factors That Regulate Glucose Production by Hepatocytes. Diabetes 68, 1156–1167, doi:10.2337/db18-1236 (2019).

40 Matsuoka, H. et al. Phosphoenolpyruvate Carboxykinase, a Key Enzyme That Controls Blood Glucose, Is a Target of Retinoic Acid Receptor-Related Orphan Receptor alpha. PLoS One 10, e0137955, doi:10.1371/journal.pone.0137955 (2015).

41 Hampson, L. J. & Agius, L. Acetylcholine exerts additive and permissive but not synergistic èects with insulin on glycogen synthesis in hepatocytes. FEBS Lett 581, 3955–3960, doi:10.1016/j.febslet.2007.07.026 (2007).

42 Gut, P. et al. Whole-organism screening for gluconeogenesis identifies activators of fasting metabolism. Nat Chem Biol 9, 97–104, doi:10.1038/nchembio.1136 (2013).

43 Sharrock, A. V. et al. NTR 2.0: a rationally engineered prodrug-converting enzyme with substantially enhanced èicacy for targeted cell ablation. Nat Methods **19**, 205–215, doi:10.1038/s41592-021-01364-4 (2022).

44 Wiles, T. J. et al. Modernized Tools for Streamlined Genetic Manipulation and Comparative Study of Wild and Diverse Proteobacterial Lineages. mBio 9, doi:10.1128/mBio.01877-18 (2018).

45 Pajak, B. et al. 2-Deoxy-d-Glucose and Its Analogs: From Diagnostic to Therapeutic Agents. Int J Mol Sci 21, doi:10.3390/ijms21010234 (2019).

46 Franza, M. et al. Zebrafish (Danio rerio) as a Model System to Investigate the Role of the Innate Immune Response in Human Infectious Diseases. Int J Mol Sci 25, doi:10.3390/ijms252212008 (2024).

47 Barton, J. R. et al. Enteroendocrine cell regulation of the gut-brain axis. Front Neurosci 17, 1272955, doi:10.3389/fnins.2023.1272955 (2023).

48 Tolhurst, G. et al. Short-chain fatty acids stimulate glucagon-like peptide-1 secretion via the G-protein-coupled receptor FFAR2. Diabetes 61, 364–371, doi:10.2337/db11-1019 (2012).

49 Meents, J. E., Ciotu, C. I. & Fischer, M. J. M. TRPA1: a molecular view. J Neurophysiol 121, 427–443, doi:10.1152/jn.00524.2018 (2019).

50 Xie, Z. et al. The gut-to-brain axis for toxin-induced defensive responses. Cell 185, 4298–4316 e4221, doi:10.1016/j.cell.2022.10.001 (2022).

51 Touhara, K. K. et al. Parasites trigger epithelial cell crosstalk to drive gut-brain signalling. Nature 653, 465–473, doi:10.1038/s41586-026-10281-5 (2026).

52 Hirabayashi, T., Nakamachi, T. & Shioda, S. Discovery of PACAP and its receptors in the brain. J Headache Pain **19**, 28, doi:10.1186/s10194-018-0855-1 (2018).

53 Rajbhandari, A. K., Barson, J. R., Gilmartin, M. R., Hammack, S. E. & Chen, B. K. The functional heterogeneity of PACAP: Stress, learning, and pathology. Neurobiol Learn Mem 203, 107792, doi:10.1016/j.nlm.2023.107792 (2023).

54 Pirone, A. et al. Pituitary adenylate cyclase-activating peptide (PACAP) immunoreactivity distribution in the small intestine of the adult New Hampshire chicken. Acta Histochem 113, 477–483, doi:10.1016/j.acthis.2010.04.008 (2011).

55 Sundler, F. et al. Pituitary adenylate cyclase activating peptide: a novel vasoactive intestinal peptide-like neuropeptide in the gut. Neuroscience 46, 439–454, doi:10.1016/0306-4522(92)90064-9 (1992).

56 Haber, A. L. et al. A single-cell survey of the small intestinal epithelium. Nature 551, 333–339, doi:10.1038/nature24489 (2017).

57 Prescott, S. L. & Liberles, S. D. Internal senses of the vagus nerve. Neuron 110, 579–599, doi:10.1016/j.neuron.2021.12.020 (2022).

58 Granton, E. et al. Biofilm exopolysaccharides alter sensory-neuron-mediated sickness during lung infection. Cell 187, 1874–1888 e1814, doi:10.1016/j.cell.2024.03.001 (2024).

59 Huerta, T. S., Tracey, K. J., Chiu, I. M. & Chang, E. H. Vagal neuroimmune circuits in inflammation and immunity. J Exp Med 223, doi:10.1084/jem.20260135 (2026).

60 McGuinness, O. P. Defective glucose homeostasis during infection. Annu Rev Nutr 25, 9–35, doi:10.1146/annurev.nutr.24.012003.132159 (2005).

61 Lang, C. H. & Dobrescu, C. Gram-negative infection increases noninsulin-mediated glucose disposal. Endocrinology 128, 645–653, doi:10.1210/endo-128-2-645 (1991).

62 Holt, R. I. G., Cockram, C. S., Ma, R. C. W. & Luk, A. O. Y. Diabetes and infection: review of the epidemiology, mechanisms and principles of treatment. Diabetologia 67, 1168–1180, doi:10.1007/s00125-024-06102-x (2024).

63 Malemnganba, T., Pandey, A. K., Kar, S. & Prajapati, V. K. Glucose Metabolism in Immune Regulation: From Mechanisms to Therapeutic Opportunities. J Cell Biochem 127, e70111, doi:10.1002/jcb.70111 (2026).

64 Britt, E. C. et al. Activation induces shift in nutrient utilization that dìerentially impacts cell functions in human neutrophils. Proc Natl Acad Sci U S A 121, e2321212121, doi:10.1073/pnas.2321212121 (2024).

65 Lane, T. A. & Lamkin, G. E. A reassessment of the energy requirements for neutrophil migration: adenosine triphosphate depletion enhances chemotaxis. Blood 64, 986–993 (1984).

66 Weisdorf, D. J., Craddock, P. R. & Jacob, H. S. Granulocytes utilize dìerent energy sources for movement and phagocytosis. Inflammation 6, 245–256, doi:10.1007/BF00916406 (1982).

67 Li, D. D. et al. Fungal sensing enhances neutrophil metabolic fitness by regulating antifungal Glut1 activity. Cell Host Microbe 30, 530–544 e536, doi:10.1016/j.chom.2022.02.017 (2022).

68 Britt, E. C. et al. Switching to the cyclic pentose phosphate pathway powers the oxidative burst in activated neutrophils. Nat Metab 4, 389–403, doi:10.1038/s42255-022-00550-8 (2022).

69 Lika, J. & Fan, J. Carbohydrate metabolism in supporting and regulating neutrophil èector functions. Curr Opin Immunol **91**, 102497, doi:10.1016/j.coi.2024.102497 (2024).

70 Sharma, K., Akre, S., Chakole, S. & Wanjari, M. B. Stress-Induced Diabetes: A Review. Cureus 14, e29142, doi:10.7759/cureus.29142 (2022).

71 Hargrove, D. M., Bagby, G. J., Lang, C. H. & Spitzer, J. J. Adrenergic blockade does not abolish elevated glucose turnover during bacterial infection. Am J Physiol 254, E16–22, doi:10.1152/ajpendo.1988.254.1.E16 (1988).

72 Matteoli, G. & Boeckxstaens, G. E. The vagal innervation of the gut and immune homeostasis. Gut 62, 1214–1222, doi:10.1136/gutjnl-2012-302550 (2013).

73 Travagli, R. A., Hermann, G. E., Browning, K. N. & Rogers, R. C. Brainstem circuits regulating gastric function. Annu Rev Physiol 68, 279–305, doi:10.1146/annurev.physiol.68.040504.094635 (2006).

74 Herman, M. A., Cruz, M. T., Sahibzada, N., Verbalis, J. & Gillis, R. A. GABA signaling in the nucleus tractus solitarius sets the level of activity in dorsal motor nucleus of the vagus cholinergic neurons in the vagovagal circuit. Am J Physiol Gastrointest Liver Physiol 296, G101–111, doi:10.1152/ajpgi.90504.2008 (2009).

75 Browning, K. N. & Travagli, R. A. Plasticity of vagal brainstem circuits in the control of gastrointestinal function. Auton Neurosci 161, 6–13, doi:10.1016/j.autneu.2010.11.001 (2011).

76 Tan, Z., Fogel, R., Jiang, C. & Zhang, X. Galanin inhibits gut-related vagal neurons in rats. J Neurophysiol 91, 2330–2343, doi:10.1152/jn.00869.2003 (2004).

77 Jèerson, K. K. What drives bacteria to produce a biofilm? FEMS Microbiol Lett 236, 163–173, doi:10.1016/j.femsle.2004.06.005 (2004).

78 Flemming, H. C. et al. Biofilms: an emergent form of bacterial life. Nat Rev Microbiol 14, 563–575, doi:10.1038/nrmicro.2016.94 (2016).

79 Melancon, E. et al. Best practices for germ-free derivation and gnotobiotic zebrafish husbandry. Methods Cell Biol 138, 61–100, doi:10.1016/bs.mcb.2016.11.005 (2017).

80 Bansal, V. & Saggau, P. Digital micromirror devices: principles and applications in imaging. Cold Spring Harb Protoc 2013, 404–411, doi:10.1101/pdb.top074302 (2013).

81 Ye, L., Robertson, M. A., Hesselson, D., Stainier, D. Y. & Anderson, R. M. Glucagon is essential for alpha cell transdìerentiation and beta cell neogenesis. Development 142, 1407–1417, doi:10.1242/dev.117911 (2015).

