## Supplementary figures and images for "A gut-brain microbial sensing pathway mediates host defense and bacterial clearance"

### Figure S1_scRNA plot Trpa1EEC_v2.jpg

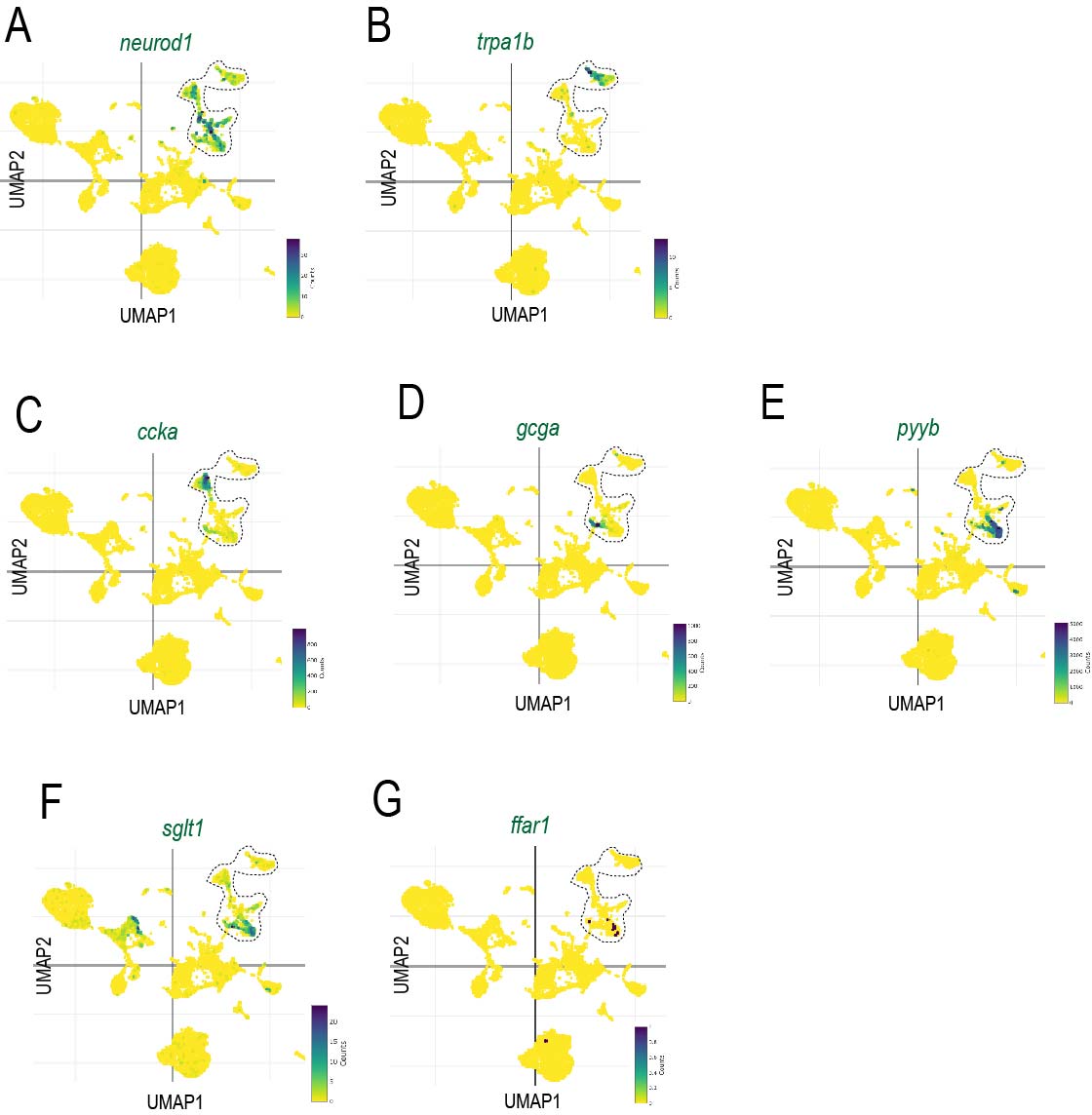

### Figure S2_EEC Gcamp-in vivo calcium.jpg

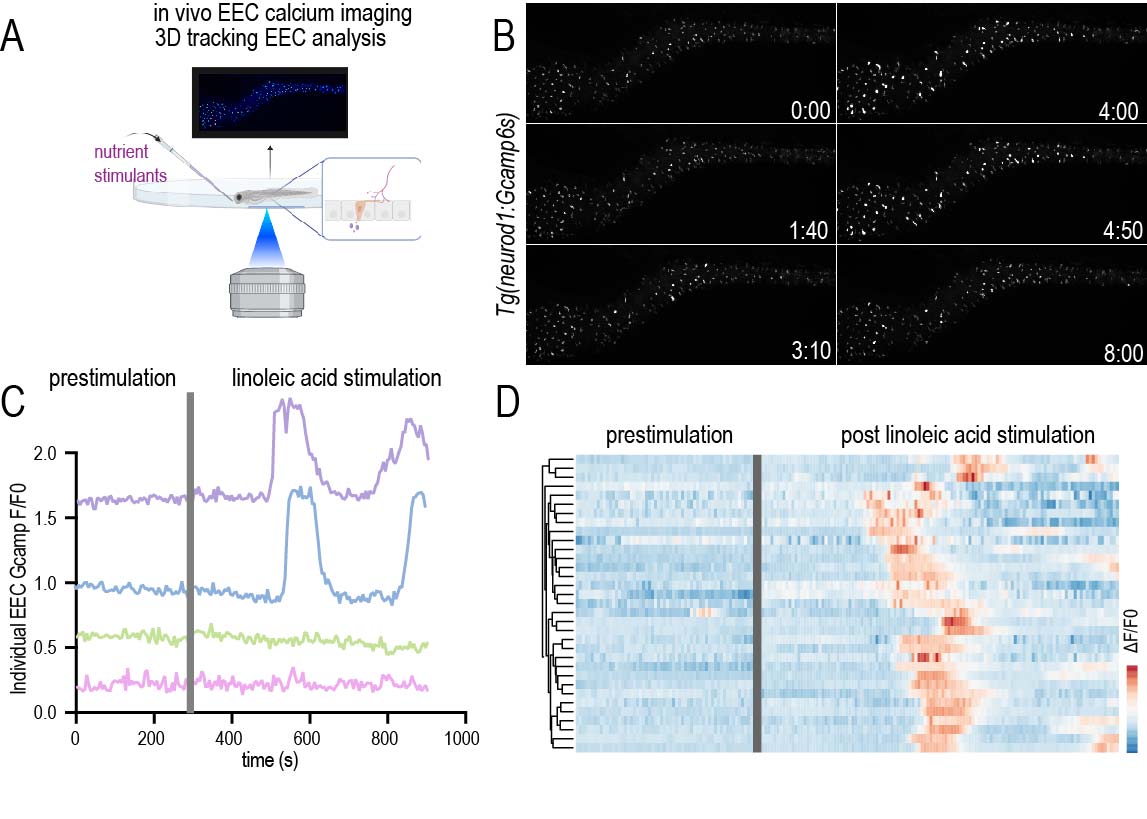

### Figure S3_light specificity.jpg

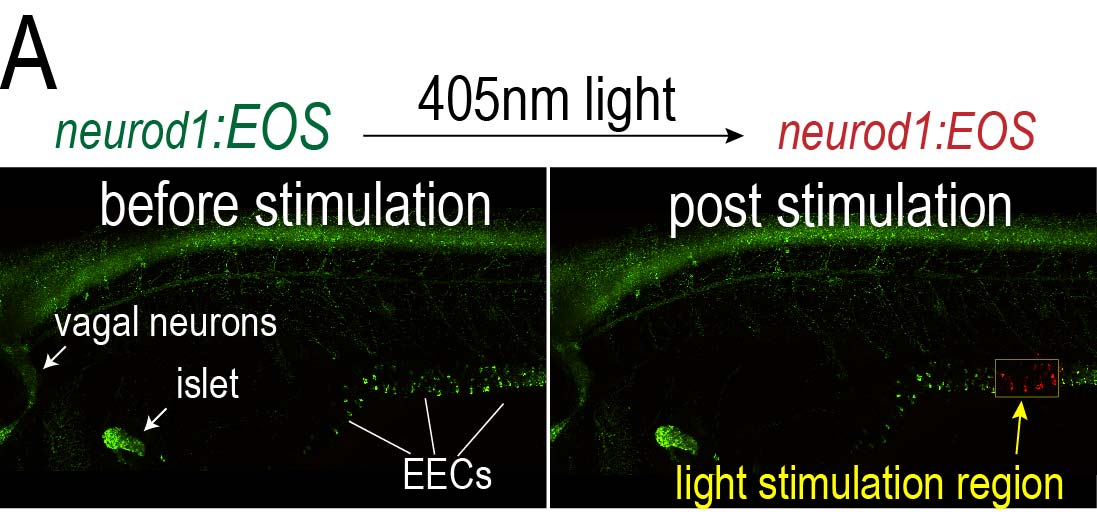

### Figure S4_Neurod1Gcamp optovin-vagal calcium_1.jpg

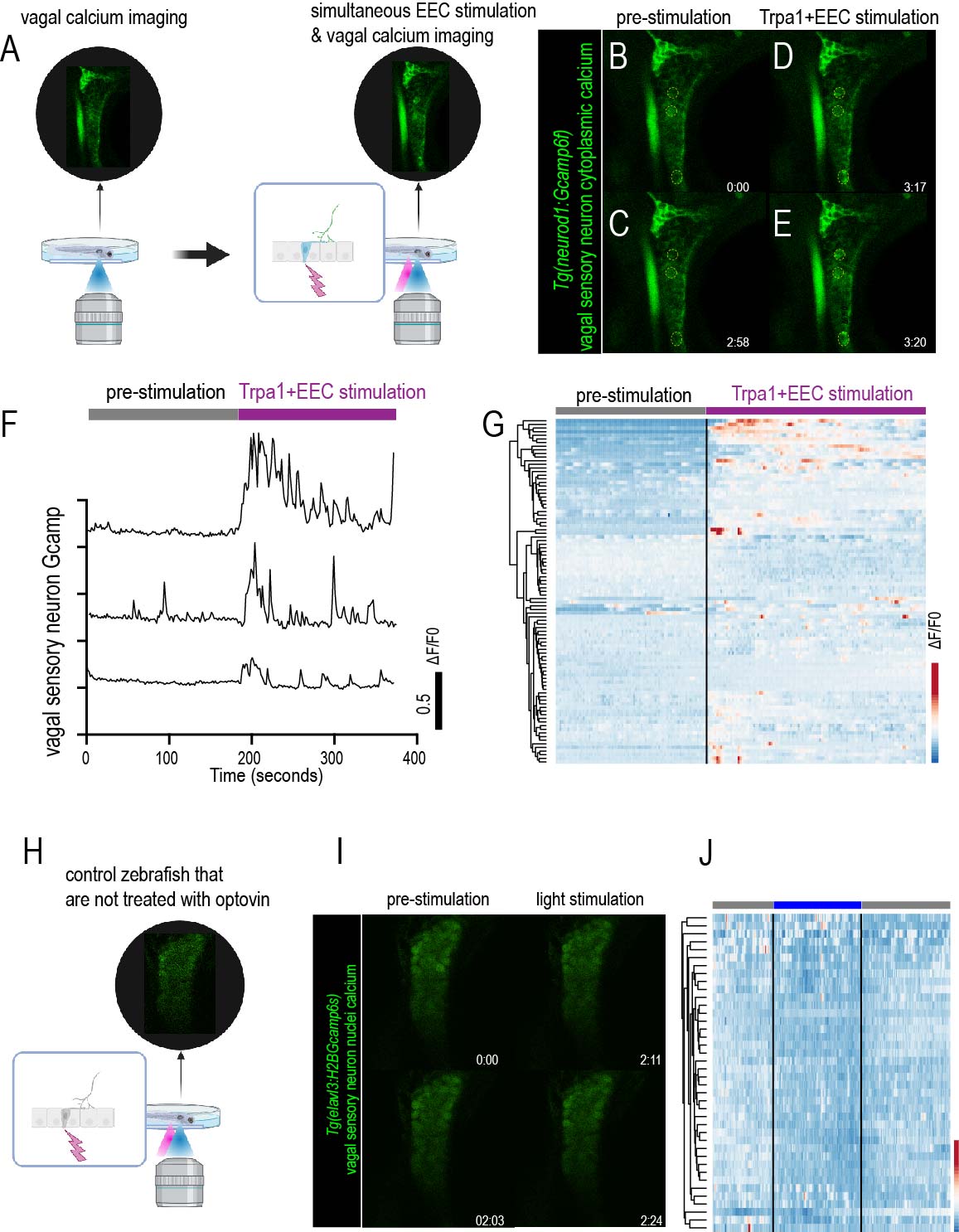

### Figure S5_glucose_water_vagal.jpg

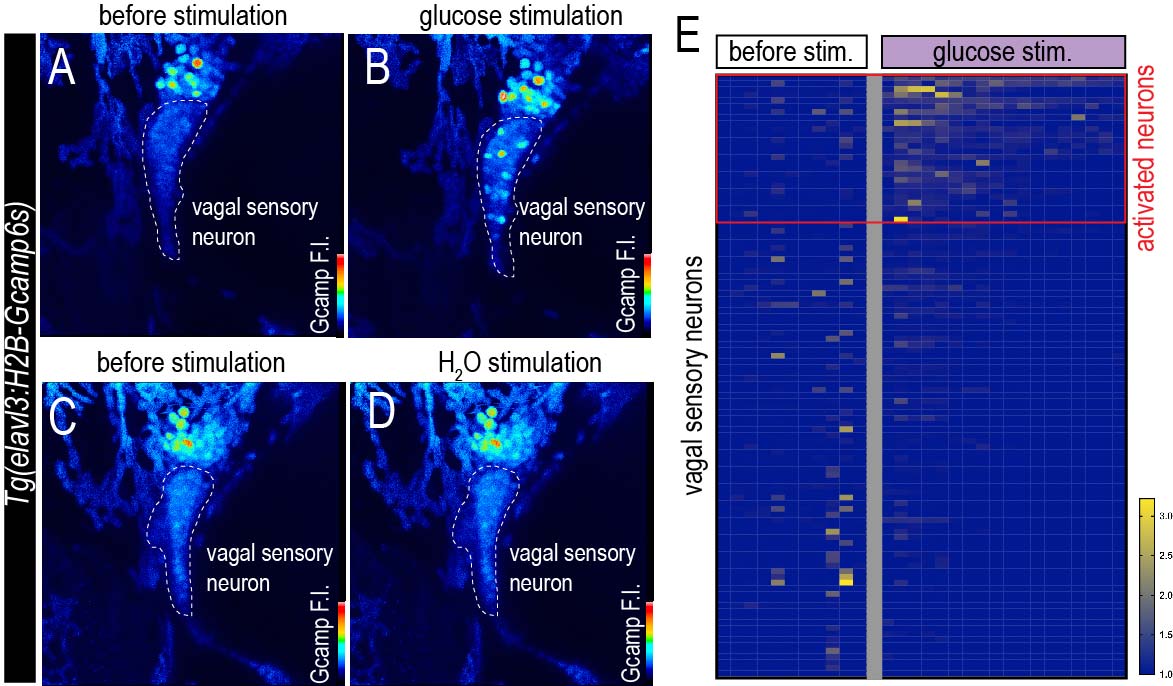

### Figure S6_htr and adcyap1r1a scRNA.jpg

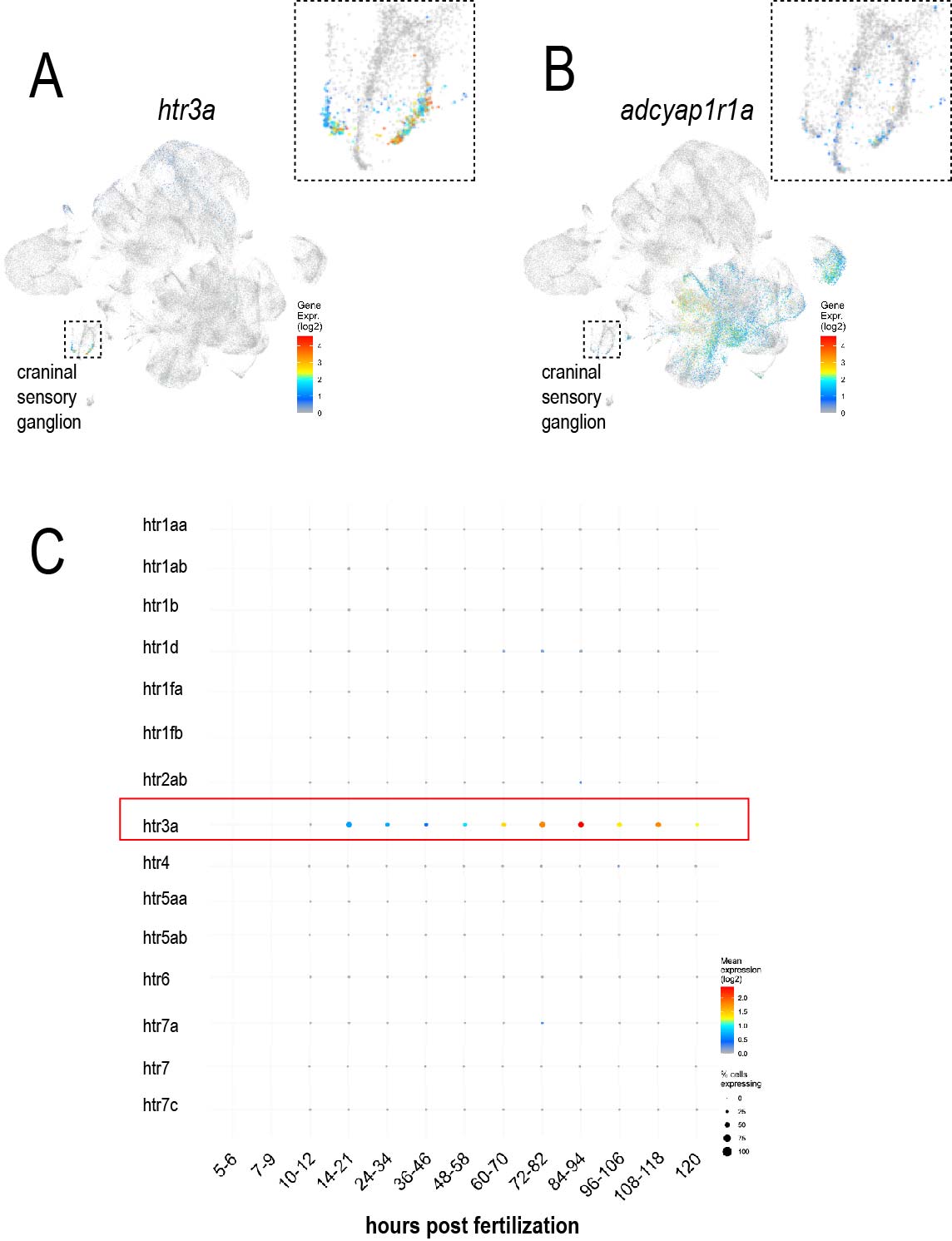

### Figure S7-adcyap1a mutant.jpg

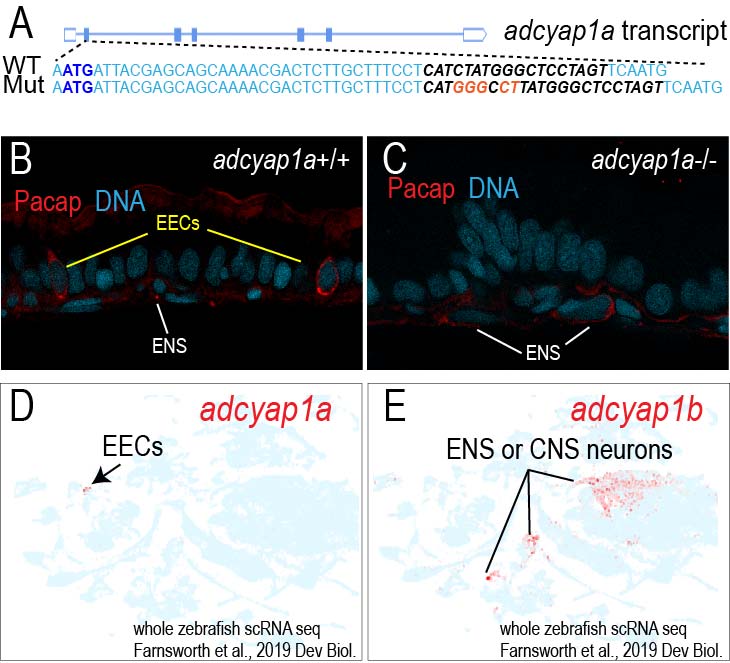

### Figure S8_chataGcamp_baseline.jpg

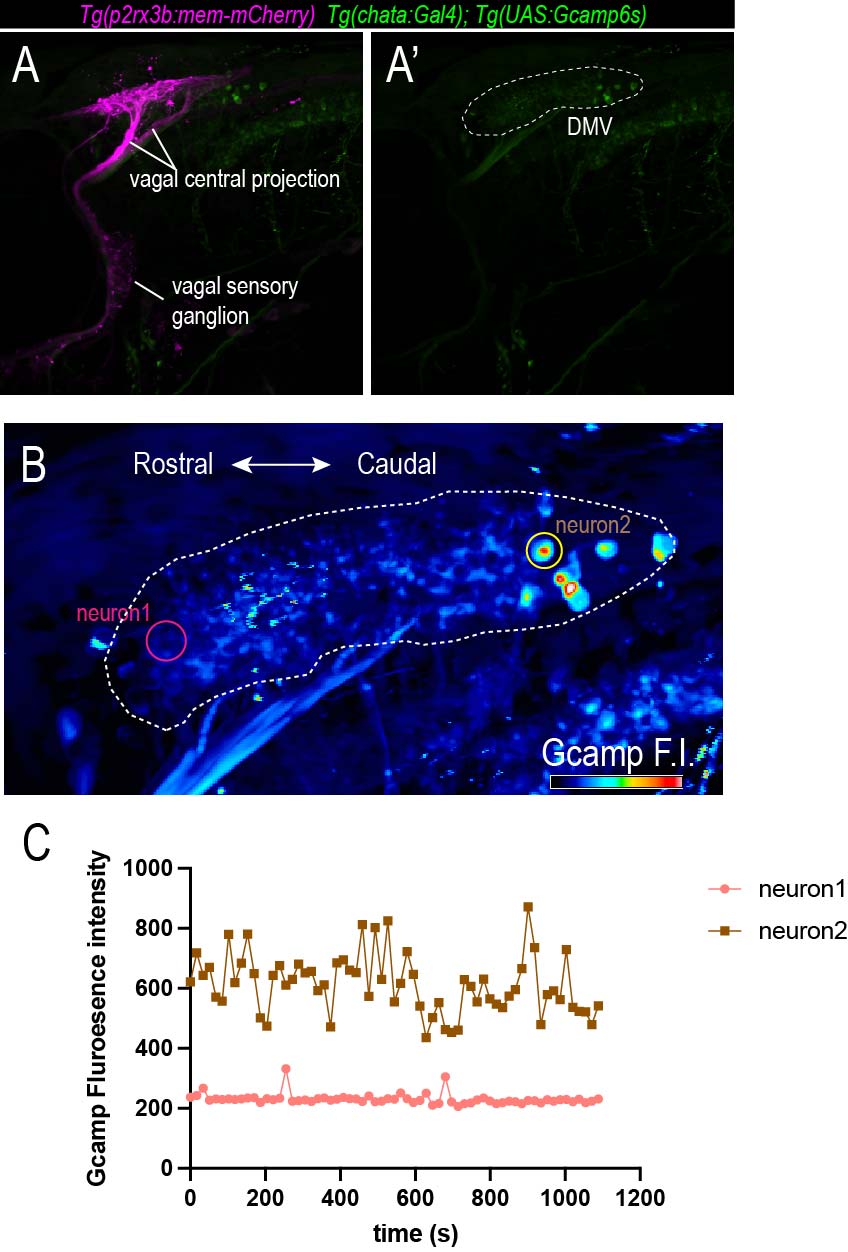

### Figure S9_Etarda cDMV pERK.jpg

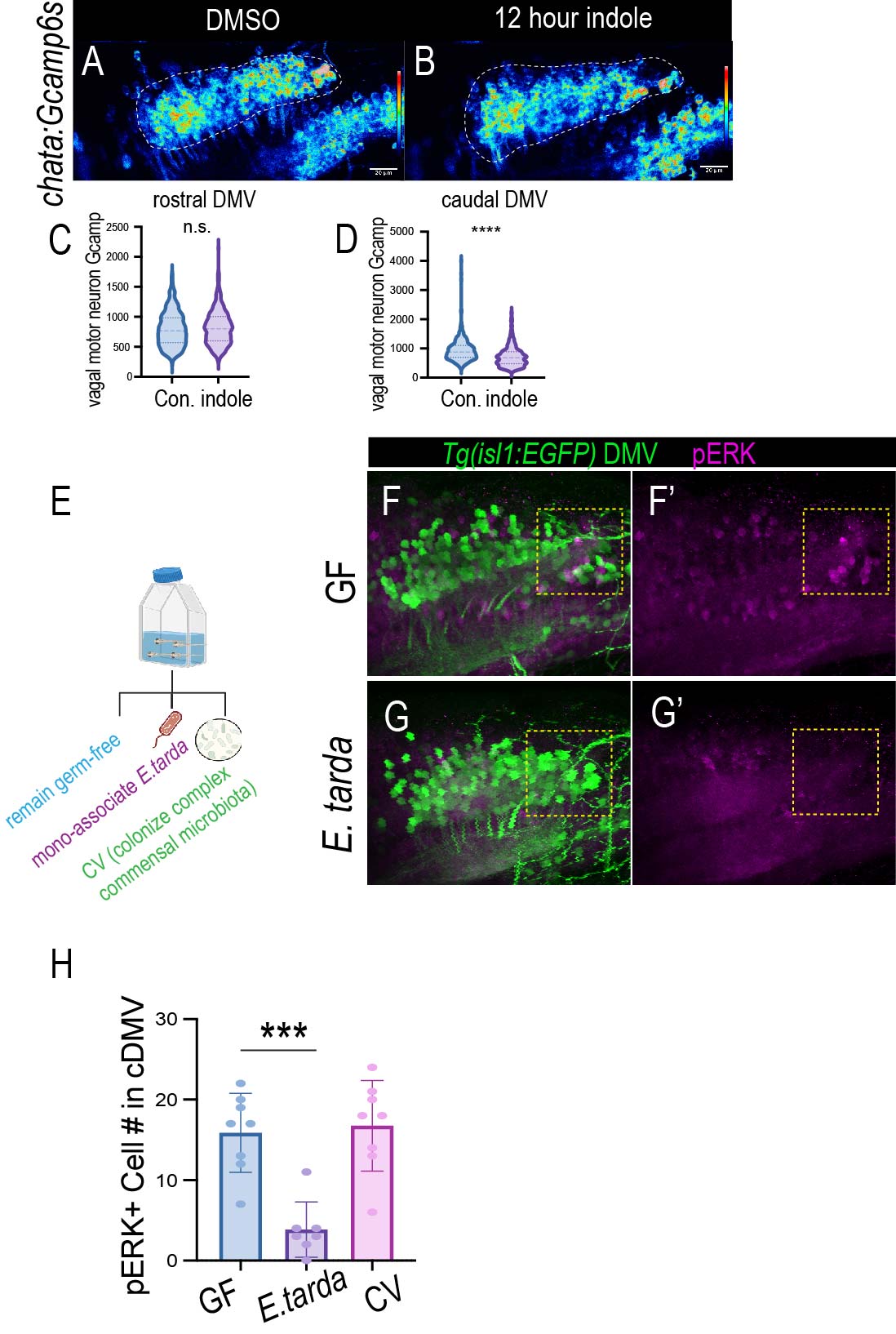

### Figure S10_adcyap1r1a and 1b liver expression.jpg

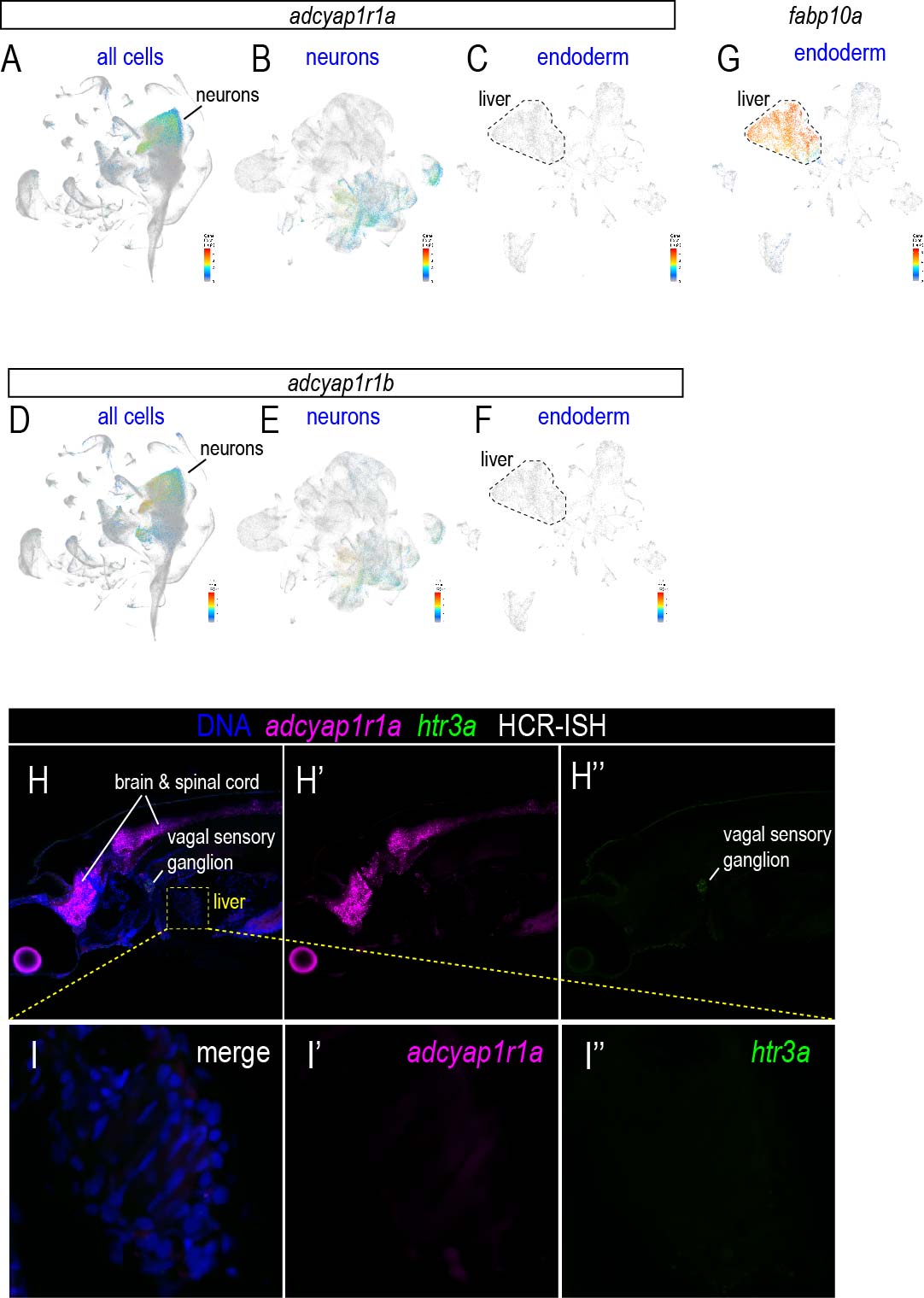

### Figure S11_adcyap1a_Etarda_v2.jpg

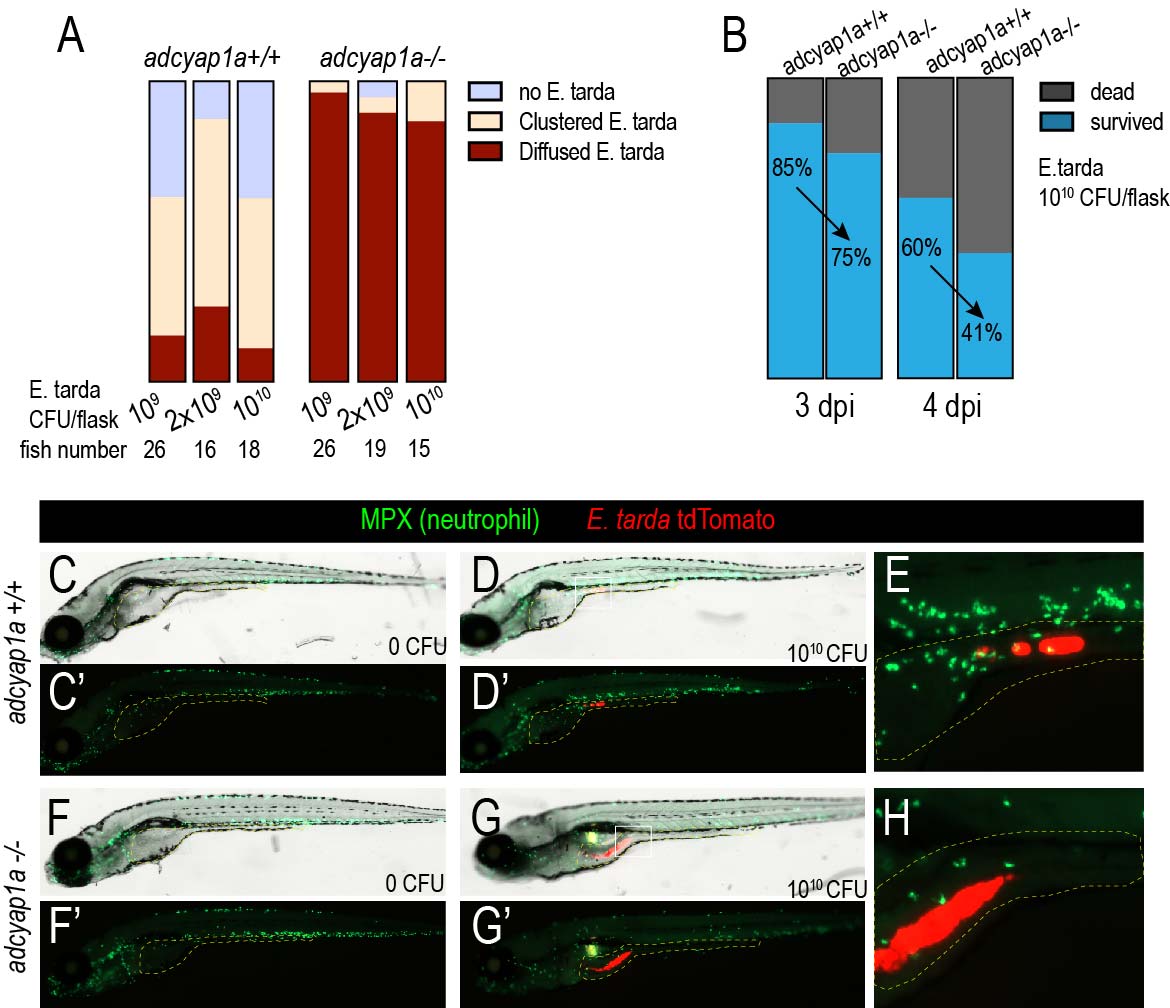
